# Lag-optimized BOLD cerebrovascular reactivity derived from breathing task data has a stronger relationship with baseline cerebral blood flow

**DOI:** 10.1101/2022.03.08.483492

**Authors:** Rachael C. Stickland, Kristina M. Zvolanek, Stefano Moia, César Caballero-Gaudes, Molly G. Bright

## Abstract

Cerebrovascular reactivity (CVR) is an important indicator of cerebrovascular health and is commonly studied with the Blood Oxygenation Level Dependent functional MRI (BOLD-fMRI) response to a vasoactive stimulus. There is theoretical and empirical evidence to suggest that baseline cerebral blood flow (CBF) modulates the BOLD signal amplitude, and that baseline CBF may influence BOLD-CVR estimates. We address how some pertinent data acquisition and modelling choices affect the relationship between baseline CBF and BOLD-CVR: whether BOLD-CVR is modelled with breathing task data or just resting-state data, and whether BOLD-CVR amplitudes are optimized for hemodynamic lag effects. For the relationship between baseline CBF and BOLD-CVR, we assessed both between-subject correlations of average GM values and within-subject spatial correlations across cortical regions. Our results suggest that a simple breathing task addition to a resting-state scan, alongside lag-optimization within BOLD-CVR modelling, can improve BOLD-CVR correlations with baseline CBF, both between- and within-subjects, likely because these CVR estimates are more physiologically accurate. We report positive coupling between baseline CBF and BOLD-CVR, both between and within subjects; the physiological explanation of this positive coupling is unclear, and future research with larger sample sizes and more tightly controlled vasoactive stimuli is needed. Understanding how baseline vascular physiology relates to dynamic cerebrovascular processes will bring further insights into what drives between and within subject participant variability in BOLD-CVR measurements and related measurements of cerebrovascular function. These insights are particularly relevant when interpreting results in populations with altered vascular and/or metabolic baselines or impaired cerebrovascular reserve.

## 1. Introduction

Cerebrovascular Reactivity (CVR), the cerebral blood flow (CBF) response to vasoactive stimuli, reflects the regulatory ability and health of the cerebrovasculature (Battisti-Charbonney et al., 2011; Meng and Gelb, 2015). Using Blood Oxygenation Level Dependent functional MRI (BOLD-fMRI) to reflect CBF changes is common for CVR mapping (Pinto et al., 2021). However, theoretical and empirical evidence shows that task-induced and resting-state BOLD amplitude changes are modulated by the vascular and metabolic baseline state (Chu et al., 2018; Griffeth et al., 2011; Kim and Ogawa, 2012; Liu, 2013; Lu et al., 2008). Multiple studies have artificially decreased or increased a subject’s resting CBF and then compared BOLD task activation amplitudes across these different states: a higher resting CBF corresponds to a lower task-based BOLD amplitude change (Brown et al., 2003; Cohen et al., 2002; Stefanovic et al., 2006; Vazquez et al., 2006). These studies primarily focus on BOLD signal changes evoked by neural activity; for BOLD-CVR effects evoked by altered blood gas levels, the literature gives less consensus on the directionality of this baseline modulation. However, there is still evidence that the baseline vascular state is coupled with CVR. For instance, positive correlations between baseline CBF (bCBF) and BOLD-CVR have been reported across individuals (Leoni et al., 2017), and studies that include both bCBF and CVR show they both decrease with age throughout certain years of adulthood (Leung et al., 2016; Lu et al., 2011). In healthy adults, there is evidence that CVR amplitudes and timings can be modulated experimentally by changing the level of baseline vasodilation or vasoconstriction (Bright et al., 2011; Halani et al., 2015).

In pathological conditions, the capacity for a vessel to dilate in response to a stimulus can be diminished due to a pre-dilated baseline, for example, in sickle cell disease bCBF is shown to be elevated and CVR diminished (Kosinski et al., 2017; Václavů et al., 2019). In steno-occlusive diseases, a vasodilatory stimulus may cause paradoxical decreases in blood flow to an area with exhausted dilatory reserve, as vascular resistance is reduced in surrounding regions (Sobczyk et al., 2014). This observation is known as the “vascular steal” phenomenon and can manifest as apparent negative CVR responses, with and without bCBF alterations (Sobczyk et al., 2014). The mechanistic relationship between bCBF and CVR in pathological cases is often not straightforward; to bring clearer interpretations to these pathological cases, a better understanding of the relationship between bCBF and BOLD-CVR in healthy populations is needed.

Variability in CVR in healthy young populations, modulated by genetic risk factors, could be a significant predicter of neurological pathology in later life (Suri et al., 2015). Therefore, understanding how baseline vascular physiology relates to dynamic cerebrovascular processes will bring further insights into what drives participant variability in BOLD-CVR measurements and related measurements of cerebrovascular function. This improved understanding will help to measure and track cerebrovascular health across the lifespan. Furthermore, BOLD-fMRI signal changes are commonly used to infer changes in neural activity, yet resting-state and task-based BOLD-fMRI amplitudes can be strongly influenced by CVR (Chen and Gauthier, 2021; Cohen et al., 2004; Liu et al., 2013b). Therefore, validity of using BOLD-fMRI as a surrogate measure of neural activity depends on the relative vascular and neural contributions to the BOLD-fMRI signal, and our ability to separate these contributions (Bright et al., 2020; Kannurpatti et al., 2010; Tsvetanov et al., 2021). These factors are particularly important to consider in clinical cohorts that present with altered vascular and/or metabolic baselines.

Here, we add to the small body of literature assessing bCBF and BOLD-CVR relationships, and address how key CVR data acquisition and modelling choices affect this relationship. Specifically, unlike most previous literature, we investigate the relationship between naturally varying bCBF and BOLD-CVR, in a healthy young population, and not in situations where bCBF has been artificially altered or in the presence or ageing or pathology. We characterize both the between-subject relationship between bCBF and CVR in gray matter (GM), and the within-subject spatial relationship of bCBF and BOLD-CVR across different cortical brain areas.

Our previous work assessed a practical modification to a typical resting-state BOLD-fMRI protocol, showing that the addition of a short breathing task to a resting-state scan facilitated CVR modelling and quantification. The inclusion of a non-invasive breathing task (breath-hold or cued deep breathing (Pinto et al., 2021)) increased the number of voxels in the brain that had a significant relationship between the partial pressure of end tidal CO_2_ (P_ET_CO_2_) and BOLD-fMRI, and improved our confidence in these voxel-wise CVR estimates. The breathing tasks evoke large, transient changes in the arterial blood CO_2_ pressure, which consequently changes blood flow; the P_ET_CO_2_ measurement is our surrogate measure of arterial blood CO_2_ changes (McSwain et al., 2010; Takano et al., 2003). If BOLD-CVR maps reflect sensible physiological variation we would hypothesize some spatially sensitive relationship with bCBF, based on previous work cited and discussed in the first paragraph of this introduction. In this paper, we assess the impact of methodological choices on between- and within-subject correlations of bCBF and BOLD-CVR. Specifically, we predict that adding breathing tasks to drive robust P_ET_CO_2_ changes will improve our detection of relationships between BOLD-CVR and bCBF. Furthermore, to achieve accurate estimates of CVR amplitude, it is important to consider spatially variable hemodynamic delays (lags) between the P_ET_CO_2_ regressor and the BOLD signal, by shifting the P_ET_CO_2_ regressor in time to optimize the full model fit (Moia et al., 2020; Stickland et al., 2021). Therefore, we also predict that lag-optimization of BOLD-CVR estimates will lead to a stronger relationship between bCBF and BOLD-CVR. Note, this manuscript focuses on the relationship between CVR maps presented previously (Stickland et al., 2021) and newly presented bCBF maps.

## 2. Methods

### 2.1. Data Collection

This study was reviewed by Northwestern University’s Institutional Review Board. All subjects gave written informed consent. See **Figure 1** for the overall protocol design. Nine healthy subjects (6 female, mean age = 26.22 ± 4.06 years) were scanned on a Siemens 3T Prisma MRI system with a 64-channel head coil. A whole brain T1-weighted EPI-navigated multi-echo MPRAGE scan was acquired, adapted from (Tisdall et al., 2016), with these parameters: 1 mm isotropic resolution, 176 sagittal slices, TR/TE1/TE2/TE3 = 2170/1.69/3.55/5.41 ms, TI = 1160 ms, FA = 7°, FOV = 256 mm^2^, Bandwidth = 650 Hz/Px, 5 minutes 12 seconds. Three T2*-weighted gradient-echo planar datasets were collected, provided by the Center for Magnetic Resonance Research (CMRR, Minnesota), with the following parameters: TR/TE = 1200/34.4 ms, FA = 62°, Multi-Band acceleration factor = 4, 60 axial slices with an ascending interleaved order, 2 mm isotropic voxels, FOV = 208 mm^2^, Phase Encoding = AP, phase partial Fourier = 7/8, Bandwidth = 2290 Hz/Px. Single band reference images were acquired to facilitate functional realignment and masking. Two of the T2*-weighted functional acquisitions included short breathing tasks preceding an 8-minute resting-state period with visual fixation, and the third acquisition only included the 8-minute resting-state period (see **Figure 1** for timings). Specifically, Breath-Hold + REST (BH + REST) included a hypercapnic breathing modulation (Bright and Murphy, 2013; Urback et al., 2017) before a rest portion, and Cued Deep Breathing + REST (CDB + REST) included a hypocapnic breathing modulation (Bright et al., 2011, 2009; Sousa et al., 2014) before a rest portion.

**Figure 1.**
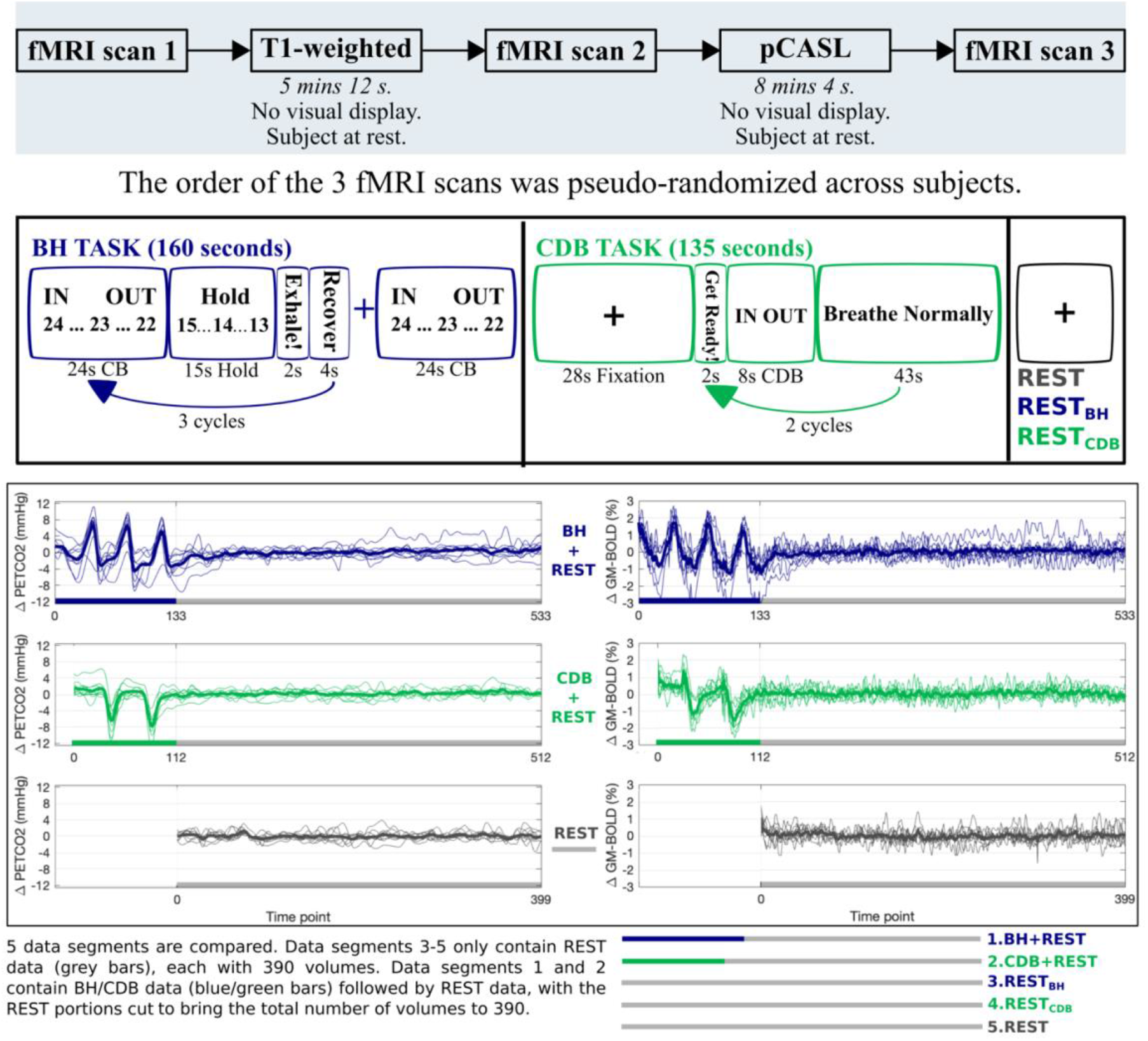
The top panel shows the study protocol. The middle panel shows the display and timings for the Breath Hold (BH) task, Cued Deep Breathing (CDB) task, and REST sections. The bottom panel displays the P_ET_CO_2_hrf traces (mmHg change from baseline) and GM-BOLD traces (% change from mean) for each fMRI scan. Each scan included 8 minutes of REST (grey), one scan preceded this with a BH task (blue) and one scan with a CDB task (green). From these three scans, five data segments of equal length were created. Thick lines represent group means, and thin lines represent each subject.

A pseudo-continuous arterial spin labelling (pCASL) dataset was also acquired whilst the subject was at rest, with sequence parameters guided by the white paper (Alsop et al., 2015a): a background-suppressed 3D GRASE read-out, 5 segments, TR/TE = 4000/19.4 ms, 4 mm isotropic voxels, 40 axial slices (interleaved, ascending), FOV = 256 mm, post label delay/Tau duration = 1800 ms, 90 mm label offset from middle slice. This sequence had 11 tag and 11 control volumes and one M0 volume, for a total scan time of approximately 7.5 minutes.

Inspired and expired CO_2_ and O_2_ pressures (in mmHg) were sampled through a nasal cannula worn by the participant, recorded at 1000Hz with LabChart software (v8.1.13, ADInstruments), connected to a ML206 Gas Analyzer and PL3508 PowerLab 8/35 (ADInstruments) and synced with volume triggers from the scanner.

### 2.2. Data Analysis

#### 2.2.1. T1-weighted image processing

The T1-weighted file was processed with FSL’s (Jenkinson et al., 2012; Li et al., 2016; Smith et al., 2004; Woolrich et al., 2009) fsl_anat function, involving brain extraction (Smith, 2002), bias field correction and tissue segmentation with FAST (Zhang et al., 2001). A GM tissue mask was subsequently created by thresholding the partial tissue volume image at 0.5.

#### 2.2.2. BOLD-CVR analysis

In brief, fMRI scans were volume registered to the same single-band reference and brain extracted. P_ET_CO_2_ values were identified, convolved with a canonical hemodynamic response function (hrf), and shifted ±15 s in 0.3 s increments, then down-sampled to the TR. Multiple linear regression was performed separately for five data segments, two of which included breathing modulations and three of which only included resting-state data. These five data segments were created from the three acquisitions as illustrated in **Figure 1,** which also displays the P_ET_CO_2_ trace and the GM average BOLD-fMRI trace for each acquisition. The regression model consisted of mean, drift terms, 6 motion parameters, and a P_ET_CO_2_ time-series. The beta-weight for the unshifted P_ET_CO_2_ regressor, scaled by the fitted mean, produced CVR maps with no lag optimization (No-Opt CVR, units: %BOLD/mmHg). For CVR maps with lag optimization (Lag-Opt CVR), the model was run for each shifted P_ET_CO_2_ regressor; parameter estimates were taken from the model with the largest full model R-squared (Moia et al., 2020; Stickland et al., 2021). For further details, refer to our previous work where we present the analysis of these CVR maps using the same subjects (Stickland et al., 2021).

#### 2.2.3. Baseline CBF (bCBF) analysis

First, volume registration was run with AFNI’s (Cox, 1996) 3dvolreg, and then the M0 volume was separated from the tag-control volumes. Brain extraction with FSL was performed on the M0 image to create a brain mask. FSL’s BASIL toolbox (Chappell et al., 2009) was used for perfusion modelling and quantification of voxel-wise cerebral blood flow in ml/100g/min. Guided by the ASL white paper (Alsop et al., 2015a) the analysis inputs included: inflow time of 3.6 seconds (post label delay of 1.8), bolus duration of 1.8 seconds, T1 of 1.3 seconds, adaptive spatial smoothing, inversion efficiency of 0.85, M0 brain masking and voxel-wise calibration with the M0 image. The fsl_anat directory was given as an input to BASIL, which performed registration to T1-weighted space.

#### 2.2.4. Transformation of atlas and CVR parameter files to T1-weighted space

In FSL, left and right hemisphere brain masks were created based on the MNI152 6th generation template brain at 2mm resolution (Grabner et al., 2006). These hemisphere masks were used alongside the Harvard Oxford Cortical Atlas (maxprob-thr25-2mm) (Desikan et al., 2006), which parcellates the cortex into 48 regions, to make an atlas with 96 cortical parcels (HarvOx_96). The fsl_anat function outputs a linear transformation matrix from T1 space to MNI space; this was inverted and the HarvOx_96 atlas file was linearly transformed to T1 space for each subject using FSL FLIRT (Jenkinson et al., 2002; Jenkinson and Smith, 2001), with nearest neighbor interpolation and 12 degrees of freedom. The single band reference image from the middle (second) fMRI scan was registered to the preprocessed T1-weighted image and the CVR maps were linearly transformed from fMRI space to T1-space for each subject. This resulted in the HarvOx_96 atlas file, CVR maps (No-Opt and Lag-Opt) and CBF maps all in T1-weighted space for each subject (**Figure 2**).

**Figure 2.**
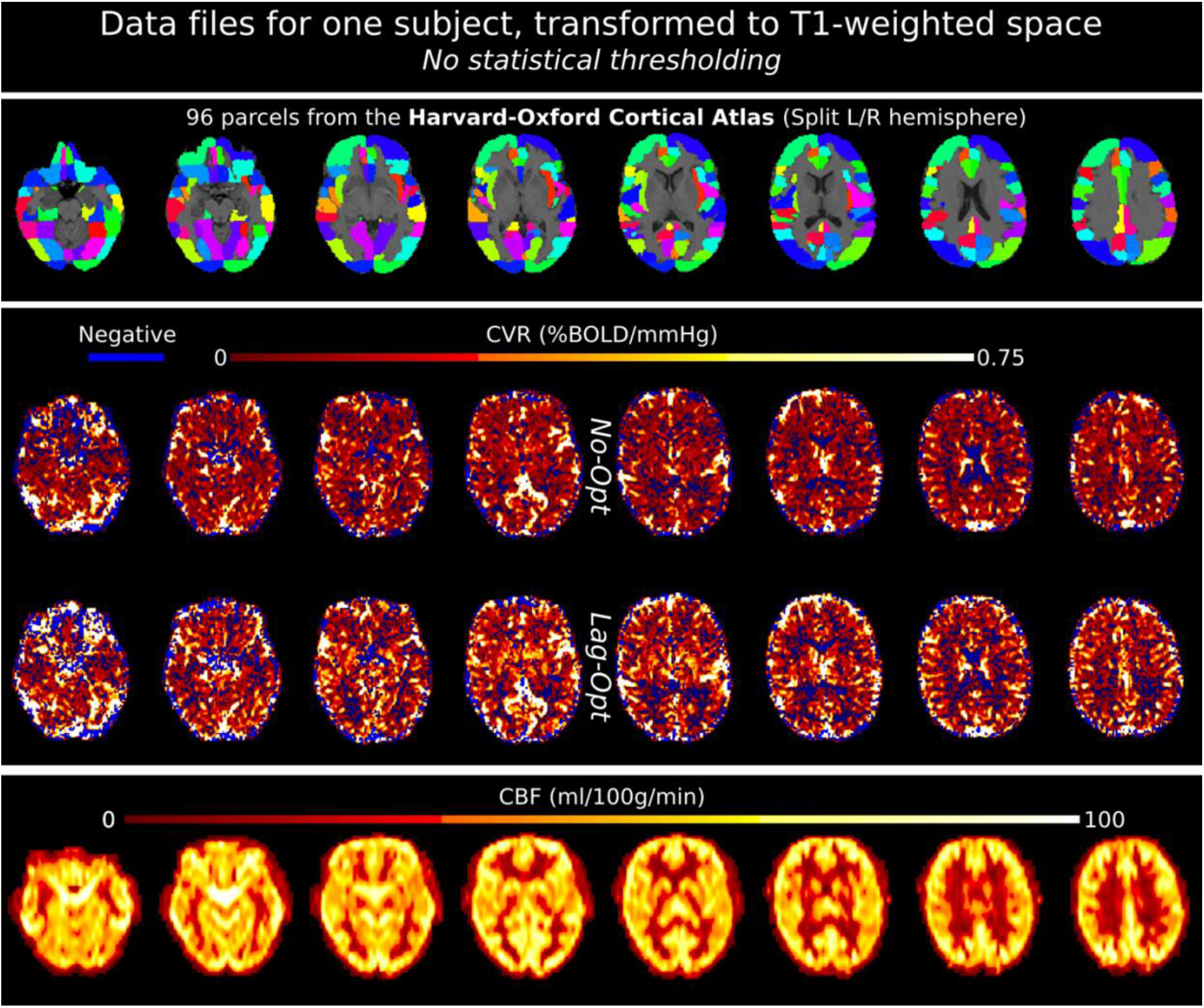
Example data files for one subject, in T1-weighted space. Top panel shows the Harvard-Oxford Cortical atlas: the initial 48 parcels were split into left and right hemisphere parcels making a total of 96. The middle panel shows CVR maps, not optimized for lag (No-Opt) and optimized for lag (Lag-Opt), for the BH+REST data segment. The bottom panel shows baseline CBF maps.

#### 2.2.5. Between-subject correlation of GM average baseline CBF and BOLD-CVR

A representative GM CVR value and GM CBF value was computed by taking a median value over voxels inside the GM mask. For each of the five data segments, and for each optimization scheme (No-Opt and Lag-Opt), Pearson’s correlation was run between these GM bCBF and BOLD-CVR values. Correlation plots and statistical outputs were created with the R packages ggplot2 (Wickham, 2016) and ggpubr (Kassambara, 2020) with the ‘ggscatter’ function. The Shapiro-Wilk test was used to ensure normality of variables.

#### 2.2.6. Within-subject spatial correlation of baseline CVR and BOLD-CVR across cortical regions

First, an average value of BOLD-CVR and bCBF within each atlas parcel was computed. Second, for each analysis combination and each subject, a spatial correlation (Spearman’s rank) between bCBF and BOLD-CVR was computed using custom MATLAB code (MathWorks, R2018b). The effect of data segment and lag optimization on bCBF-CVR spatial correlation values was tested with a repeated measures ANOVA, run with the R package permuco (Frossard and Renaud, 2019) with the ‘aovperm’ function. Null distributions were created via 100,000 permutations of the original data, which therefore do not depend on Gaussian and sphericity assumptions. When investigating simple main effects (‘emmeans’ package) and performing multiple comparisons, p-values were false discovery rate (FDR) corrected, and then compared against an alpha of 0.05 to determine significance.

#### 2.2.7. Within-subject correlation of resting-state metrics and bCBF across cortical regions

Results from our previous work (Stickland et al., 2021) and from the analyses described in sections 2.2.5 and 2.2.6 show the difficulty of estimating BOLD-CVR with resting-state data, within a lagged-GLM framework using a P_ET_CO_2_ regressor. However, literature suggests that metrics derived directly from BOLD resting-state fluctuations may capture CVR effects (Chen and Gauthier, 2021; Pinto et al., 2021), albeit not in quantitative units, making comparisons between subjects or between time-points more limited. We performed a simple exploratory analysis looking at the spatial relationship between resting-state metrics and bCBF. Minimally pre-processed BOLD-fMRI data (brain extraction, volume registration) from the three resting-state data segments (REST, REST_BH_ and REST_CDB_) were input to AFNI’s 3dRSFC function (including quadratic detrending of input data, bandpass filtering between 0.01 and 0.1 Hz and spatial smoothing with a 4mm full width half maximum (FWHM) kernel). This function computed maps of resting-state fluctuation amplitude normalized to the mean (mRSFA), amplitude of low-frequency fluctuation (ALFF) (Zang et al., 2007) and fractional ALFF (fALFF) (Zou et al., 2008). ALFF and RSFA (Kannurpatti and Biswal, 2008) are similar metrics; the normalized version of RSFA (mRSFA) better distinguishes this metric from ALFF (Golestani et al., 2016; Liu et al., 2013a). The same spatial correlations with bCBF were performed for each subject as explained in Section 2.2.6.

## 3. RESULTS

### 3.1. Between-subject correlation of GM average baseline CBF and BOLD-CVR

**Figure 3** shows the relationship between GM bCBF and GM BOLD-CVR across subjects, for each data segment and lag optimization scheme. After lag optimization (and with influential points removed), all correlations between bCBF and CVR are positive: a higher bCBF tends to co-occur with a higher CVR across subjects. Importantly, the only significant correlations with bCBF were found with lag optimized CVR values derived from data segments with breathing tasks included (i.e., BH+REST and CBD+REST). However, no correlations were found to be statistically significant after FDR correction across the 10 tests. These results are consistent across different GM masking options (Supplementary Table 1).

**Figure 3.**
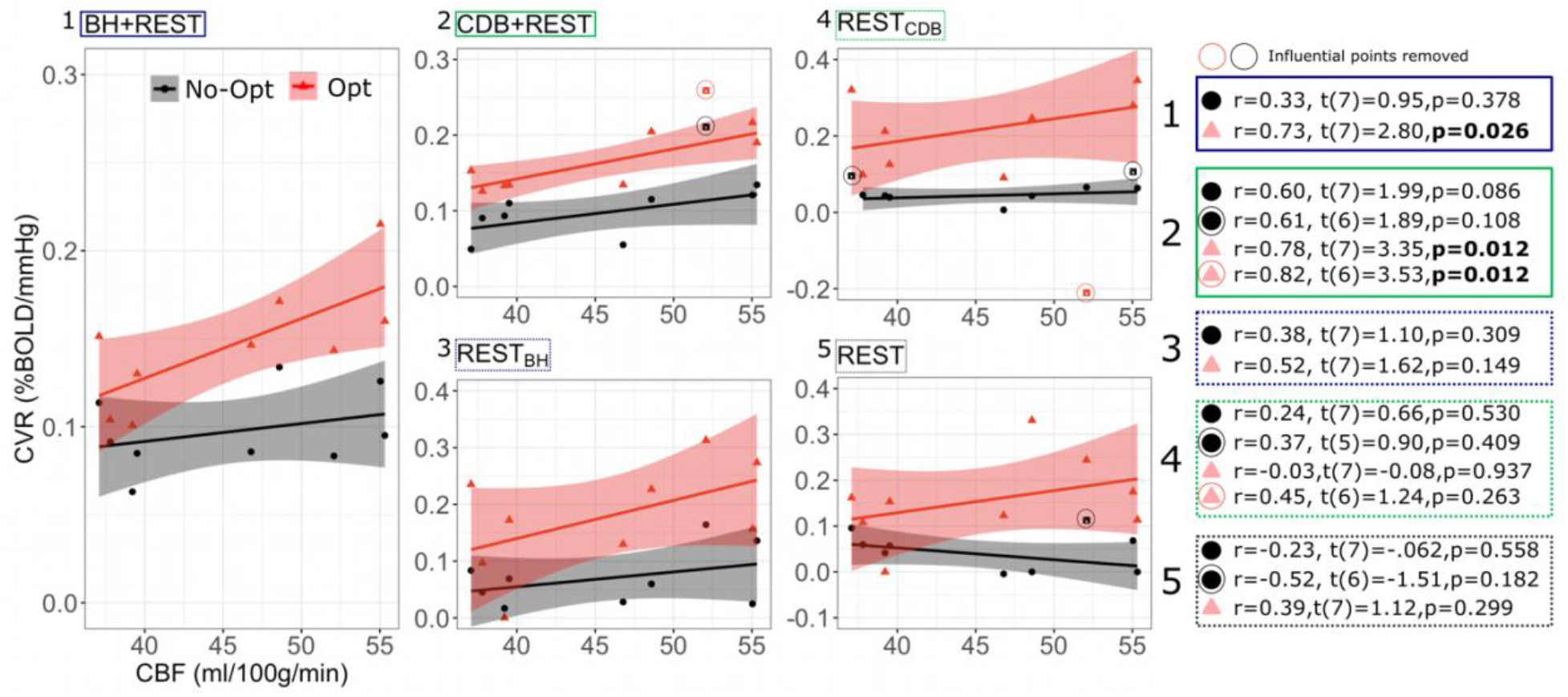
Between-subject Pearson correlations between GM CBF values and GM BOLD-CVR values, shown for each data segment (1-5) and for CVR values with no lag optimization (No-Opt, black circles) and CVR values with lag optimization (Opt, pink triangles). The CBF and CVR values are medians over a GM mask and each dot represents one participant. On the right, statistical tests are shown with and without influential points removed. Influential points were classed as points with a Cook’s distance over 4/n, with n being the number of subjects. P-values are not corrected for multiple comparisons. The gray shaded regions around the fit lines indicate the 95% confidence interval of the correlation coefficient (fit line and confidence interval estimation did not include influential points, but influential points are still plotted alongside).

### 3.2. Within-subject spatial correlation of baseline CBF and BOLD-CVR across cortical regions

**Figure 4** illustrates the spatial correlation, for each subject, across the 96 cortical parcels of the atlas. Single subject correlations that are significant (outside the pink shaded box in panel B) are mostly positive. Panel C of Figure 4 shows how bCBF and BOLD-CVR correlations changed due to lag optimization. This change was most consistent for BH+REST data, with 8/9 subjects showing an increased (positive) correlation after lag optimization. For the other segments, changes with lag optimization are more variable, and there are more negative correlations. The repeated measures ANOVA showed no significant interaction effect (F(4,32)=0.77, p=0.5552) and two significant main effects: correlation values were significantly higher for Lag-Opt versus No-Opt (F(1,8)=6.09, p=0.0385) and there was a significant effect of data segment on correlation values (F(4,32)=3.84, p=0.0117). BH+REST showed significantly higher values than REST_CDB_ (see Supplementary Table 2 for all pairwise comparisons).

**Figure 4.**
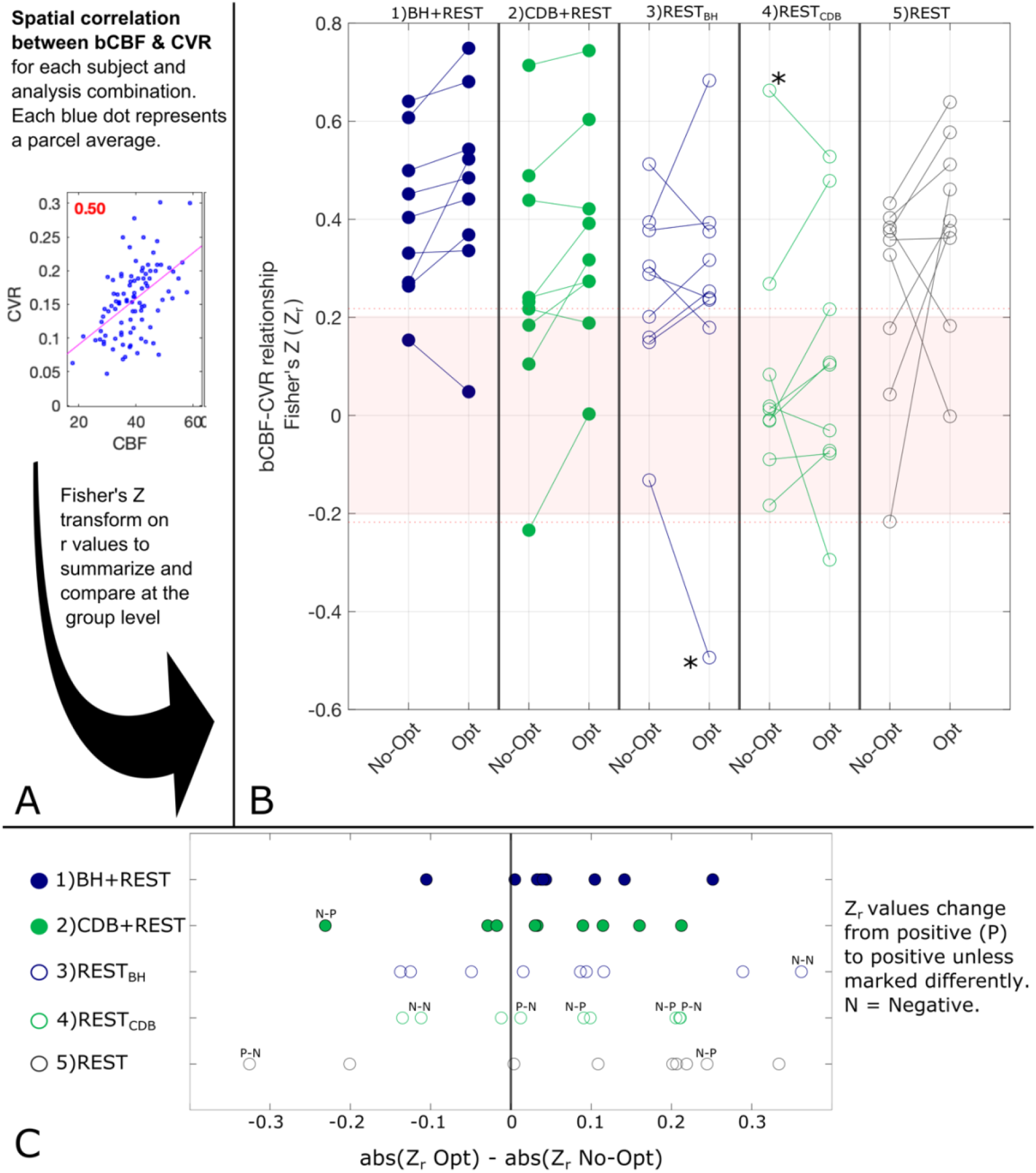
The association between bCBF and CVR values in 96 cortical parcels was computed for each subject and analysis combination (panel A). A Fisher’s Z transformation is applied to the correlation values to allow group statistical testing; panel B visualizes these transformed correlations for the five segments (columns), both for CVR values with no lag optimization (No-Opt) and CVR values with lag optimization (Opt). The shaded pink box in panel B represents correlations that were **not** significant at the single-subject level, based on 96 datapoints and a critical value of 1.96 (p<0.05, two-tailed). When generating these correlation values for each subject, outlier parcels (based on Cook’s distance) were not included. The number of parcels removed from each correlation analysis averaged 5.3 out of 96 (see Supplementary Table 3). As removing parcels reduces the degrees of freedom, the dotted pink line shows the adjusted critical value (maximum across participants and data segments shown for reference). Panel C shows the change in absolute correlation strength due to lag optimization of the CVR values.

Two extreme outlier points were identified (greater than 3 x IQR below the first quartile or above the third), indicated by an asterisk in **Figure 4**. When removing these two subjects the same results were found: no significant interaction effect (F(2,24)=0.30, p=0.8732), and significant main effects of optimization scheme (F(1,6)=8.34, p=0.02682) and data segment (F(2,24)=10.83, p=0.00006). When replacing these outlier values with the mean value for that cell of the design, the same results were also found: no significant interaction (F(4,32)=0.46, p=0.7661), and significant main effects of optimization scheme (F(1,8)=10.67, p=0.0111) and data segment (F(4,32)=7.40, p=0.00031). Multiple ANOVAs were explored due to the challenges with removing outliers, particularly for within-subject designs.

### 3.3. Within-subject correlation of resting-state metrics and bCBF across cortical regions

**Figure 5** illustrates that in the REST data segment, all single subject spatial correlations between bCBF and ALFF, fALFF and mRSFA were positive and significant at the single subject level, with fALFF showing slightly higher correlation strength with bCBF compared to ALFF and mRSFA. ALFF and mRSFA showed extremely similar maps, and therefore also similar correlation values with bCBF, which is logical because a correlation analysis is not sensitive to differences in absolute scale. A similar pattern of results was also seen for the REST_BH_ and REST_CDB_ data segments (Supplementary Figure 1). It is important to note that we generated outputs with and without smoothing (Supplementary Figure 1); the correlations are notably different for ALFF and mRSFA, showing much lower correlation values without smoothing, which are no longer significant at the single-subject level for the majority of subjects. Smoothing is a typical processing step in resting-state fMRI analyses, yet the appropriate kernel size is hard to determine. Using a Gaussian kernel size of two or three times the voxel size is a common recommendation (Alahmadi, 2021; Cumming et al., 2012; Liu et al., 2017; Pajula and Tohka, 2014); thus, we smoothed with FWHM size of 4mm, double our voxel size.

**Figure 5.**
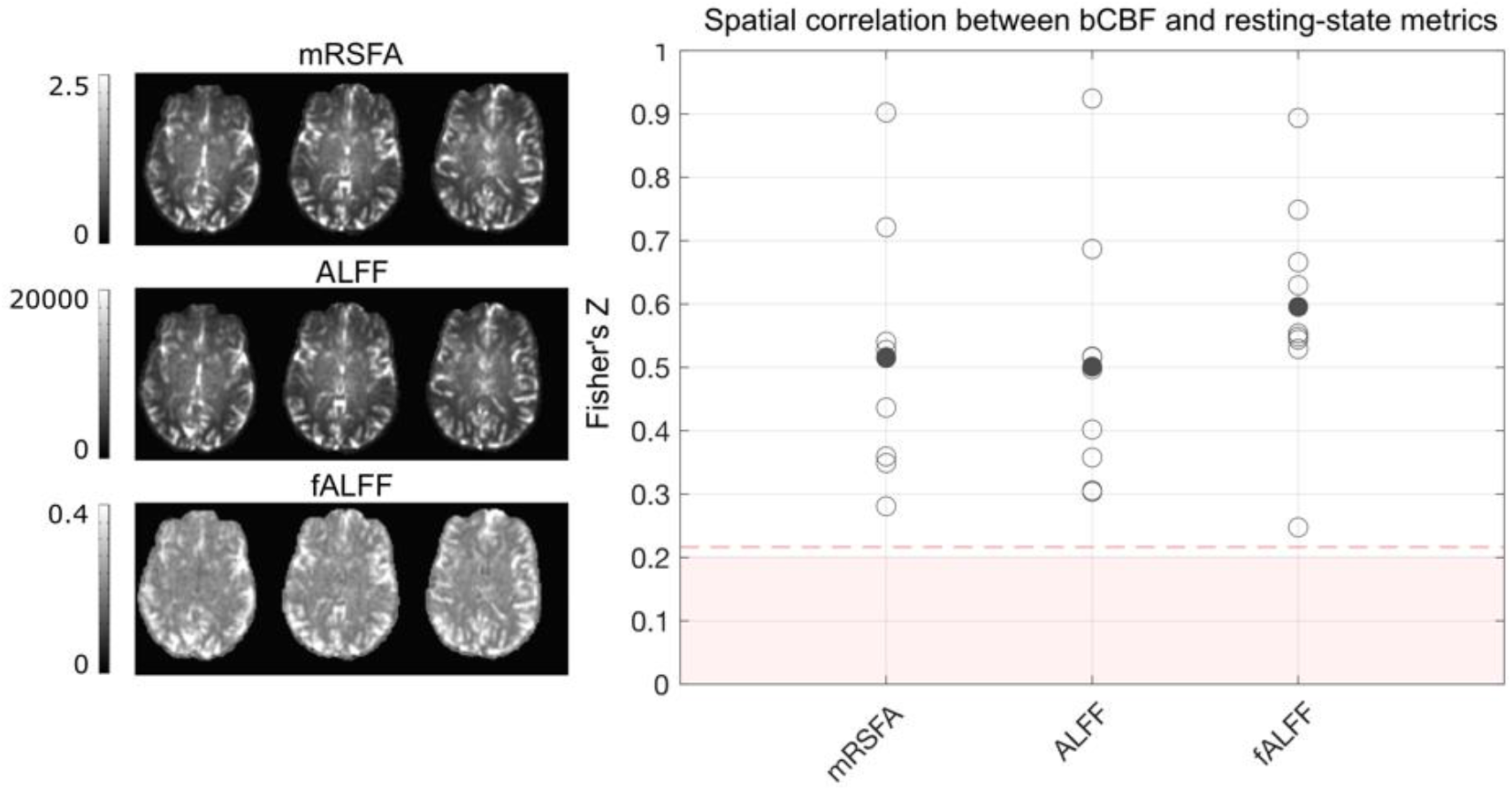
The association between bCBF and each resting-state metric, in 96 cortical parcels, was computed for each subject, using data from the REST data segment. Maps from one example subject are displayed. A Fisher’s Z transformation is applied to the correlation values before averaging across subjects. The shaded pink box represents correlation values that would **not be** significant at the single-subject level, based on 96 datapoints and a critical value of 1.96 (p<0.05, two-tailed). When generating these correlation values for each subject, outlier parcels (based on Cook’s distance) were not included. As removing parcels reduces the degrees of freedom, the dotted pink line shows the adjusted critical value (maximum across participants and resting-state metric shown for reference).

## 4. Discussion

We assessed the impact of methodological factors on the correlation between bCBF and BOLD-CVR. We present complimentary results to those in our previous manuscript (Stickland et al., 2021), suggesting that adding a simple breathing task to a resting-state BOLD fMRI scan, alongside lag-optimization of a P_ET_CO_2_ regressor within CVR modelling, improves the relationship between BOLD-CVR and baseline CBF. We also report exploratory results of positive spatial correlations between bCBF and ALFF, mRSFA and fALFF, for each of the three resting-state segments. We first discuss the impact of these methodological factors in more detail (Section 4.1), followed by exploring reasons for the observed positive relationship between bCBF and BOLD-CVR and the resting-state metrics (Section 4.2), and then consider the relevance and implications for fMRI research (Section 4.3).

### 4.1. The impact of methodological factors on the correlation between bCBF and BOLD-CVR

For variability between-subjects, a significant positive correlation between an average GM BOLD-CVR value and an average GM bCBF value was only found for BOLD-CVR data that included a breathing task and when analysis included optimization for hemodynamic lag effects. With influential points removed, the strength of the correlation always increased after lag-optimization, for all comparisons. For within-subject spatial correlations between bCBF and BOLD-CVR in cortical parcels, the bCBF and BOLD-CVR relationship was mostly positive (the negative correlations were only seen in resting-state data with no breathing tasks or in results without lag-optimization). At the group level, spatial correlations were significantly higher after lag-optimization of the CVR values. This suggests that these two methodological choices, the inclusion of breathing data and the implementation of lag-optimization, result in more physiologically accurate CVR estimates.

We have shown previously (Stickland et al., 2021) that adding breathing tasks to resting state fMRI paradigms makes CVR mapping more robust: if BOLD signal changes related to P_ET_CO_2_ are small in amplitude, as may occur if natural fluctuations in P_ET_CO_2_ are small during resting-state paradigms, it can be hard to distinguish them from other physiological, artefactual, or neuronally-driven fluctuations that occur at the same low frequencies. Our observations in this work suggest that BOLD-CVR estimates derived from resting-state data also show a much more variable relationship with bCBF. The timing of the blood flow response to a change in the pressure of arterial CO_2_ (as measured here with end-tidal CO_2_ recordings) will not be the same for each equipment setup, each participant, and each area of the brain due to methodological and physiological factors. As well as the data type to be modelled (resting state only vs. added breathing task), accounting for voxel-wise hemodynamic timings when estimating BOLD-CVR seems to lead to more physiologically grounded CVR values.

It is not clear why the REST_CDB_ segment had significantly lower spatial correlations between bCBF and BOLD-CVR than the other two REST segments, irrespective of lag-optimization. It could possibly be due to residual vascular after-effects following the hypocapnic breathing task stimulus (Bright et al., 2011). The CDB task had fewer visual instructions than the BH task, so if *neurovascular* after-effects played a role, we would expect this to be more relevant for the REST_BH_ data segment as opposed to REST_CDB_. Furthermore, the REST_CDB_ data segment that was analyzed was preceded by 55 seconds of the subject breathing normally (the last 43 seconds of the ‘breathe normally’ task cue, see Figure 1, and the first 12 seconds of the REST_CDB_ section not analyzed during CVR modelling). We previously found no clear evidence for obvious differences in common resting-state metrics (RSFA, ALFF, fALFF, LFCD) between all three resting-state data segments (Stickland et al., 2021), though further investigation of neural and vascular after-effects of breathing tasks may be warranted.

### 4.2. A positive correlation between baseline CBF and BOLD-CVR

There are many studies in the literature including both bCBF and BOLD-CVR values for healthy young populations, but few studies that directly assess the covariance of these values. In conflict with our results, a PET study of healthy subjects, of similar age to this study, showed a significant *negative* correlation between baseline CBF and CBF-CVR (with hypocapnia and hypercapnia) in multiple brain regions (Ito et al., 2008). This difference could be driven by the fact they assessed CBF-CVR, as opposed to BOLD-CVR, which we discuss further in a subsequent section. A recent study (Chen et al., 2021) with a sample of similar age to ours investigated sex differences in bCBF and BOLD-CVR, showing that female participants had higher bCBF (consistent with what we see in Supplementary Figure 2) but no difference in BOLD-CVR. However, they also reported a *negative* correlation between bCBF and BOLD-CVR, using average values from four different brain lobes (frontal, temporal, parietal, and occipital). They concluded that low and high autoregulation efficiencies exist at high and low bCBF values, respectively. This is again contrary to our results, where we see positive correlations between these metrics. There are key methodological and analytical differences between this study and ours: average values were derived from 4 brain lobes compared to the 96 cortical parcels in this study; they did not record P_ET_CO_2_ values, precluding characterization of BOLD-CVR in units of %BOLD/mmHg; the BOLD response to the breath-hold was modelled with the convolved task design and hemodynamic timings were not corrected on a regional level; CBF and BOLD fMRI data were smoothed with an 8mm FWHM Gaussian kernel whereas we did not apply any spatial smoothing except the adaptive spatial smoothing applied with FSL BASIL’s toolbox. Finally, the focus of their study was to compare sex differences in bCBF and BOLD-CVR; their CBF and CVR correlations are displayed as group averages with large standard deviations, and it is therefore hard to distinguish the contribution of individual variability to this main result.

#### 4.2.1. Ageing effects

Previous publications reporting or commenting on a *positive* correlation between bCBF and BOLD-CVR across subjects often explain this in the context of cerebrovascular aging effects: in elderly adults, a smaller capacity for vasodilation is often seen alongside lower bCBF (Leoni et al., 2017; Lu et al., 2011; Marstrand et al., 2002; Tsvetanov et al., 2021). Developmental age is an important factor to consider; one study examined how BOLD-CVR and bCBF correlated with age in a sample ranging from 9-30 years (Leung et al., 2016). BOLD-CVR and bCBF only changed in the same way with age after 14.7 years (both decreased with age), whereas between 9 and 14.7 years bCBF decreased with age and CVR *increased* with age, resulting in a negative relationship between these parameters. We cannot rule out age effects as a potential causal explanation for the bCBF-CVR relationships in our data, and we do see negative trends with age and bCBF and negative trends with age and BOLD-CVR that agree with the literature (Supplementary Figure 2). However, it is challenging to interpret our findings within the framing of ageing effects or cerebrovascular impairment given our small sample of healthy younger adults with a small age range (23-35 years), and this was not the focus of the study. Therefore, alternative interpretations separate from ageing effects are explored next.

#### 4.2.2. Neurovascular and metabolic individual differences

Under the assumption that our vasoactive breathing task stimuli are isometabolic, the BOLD-CVR response should reflect the CBF-CVR response (Zhou et al., 2015). However, there are many physiological factors that can contribute to a BOLD signal change other than a CBF change (Kim and Ogawa, 2012), and the assumption of an isometabolic state under hypercapnia may not always be valid (Deckers et al., 2021; Driver et al., 2017; Xu et al., 2011; Yablonskiy, 2011). Furthermore, BOLD-CVR and CBF-CVR may have differing dependencies on the baseline condition (Halani et al., 2015; Stefanovic et al., 2006) and on the type of vascular stimulus (Halani et al., 2015). It is therefore possible that individual variability in BOLD-CBF coupling during breathing tasks could influence our results.

Healthy individual differences in the functioning of the neurovascular unit (Schaeffer and Iadecola, 2021) could also affect both bCBF and BOLD-CVR in similar ways, leading to their positive coupling. To contextualize our results within commonly derived resting-state BOLD metrics, separate from our lagged-GLM framework, we performed an exploratory analysis to assess the relationship between resting-state metrics and bCBF. We found weak to moderate positive spatial correlations with bCBF for both ALFF and mRSFA (these are directly proportional measures). The contribution of neural or vascular factors driving voxel-wise ALFF/mRSFA values is not clear, and they have been interpreted and applied as both neural and vascular in the literature (Chen and Gauthier, 2021). Interestingly, fALFF had a stronger positive correlation with bCBF in our data compared to ALFF/mRSFA (regardless of smoothing, Supplementary Figure 1), and fALFF is thought to be more neuronally specific than ALFF. A recent paper discusses and investigates the complexities of these physiological interpretations (Deng et al., 2022), and their results suggest ALFF is more influenced by venous vascular and microvascular density (Vigneau-Roy et al., 2014), and fALFF is reflective of underlying metabolic demand. Therefore, the positive spatial correlation between fALFF and bCBF could reflect more metabolically active regions at rest requiring more blood flow. The degree of coupling between resting-state BOLD and CBF signals also relates to the macrovascular volume fraction (i.e. BOLD and CBF are less coupled in voxels near larger vessels) (Tak et al., 2014). It is therefore possible that the positive coupling of bCBF with BOLD-CVR, ALFF/mRSFA and fALFF could be caused by overlapping, yet distinct physiological drivers (i.e., BOLD driven effects, CVR driven, metabolism driven).

#### 4.2.3. Non-linear relationship between CBF and P_ET_CO_2_

This positive bCBF and BOLD-CVR coupling could reflect the non-linear relationship between CBF and P_ET_CO_2_: the sigmoidal dose-response curve combined with lowered baseline P_ET_CO_2_ levels associated with paced breathing between breath-holds, or due to sustained effects of hyperventilation in the CDB task, may strongly influence the magnitude of CBF change during breathing tasks (see, for example, Figure 7 in (Tancredi and Hoge, 2013)). However, there is evidence that CVR studies can still assume to operate in the linear portion of the dose-response curve if baseline P_ET_CO_2_ values are kept within 30 – 45 mmHg (Tancredi and Hoge, 2013), as is the case with our data (Supplementary Table 4, Part B). Nevertheless, CVR studies using breathing tasks should take care to avoid causing undue hypocapnia during baseline periods, whilst attempting to achieve stability with paced breathing, as this could bias CVR measurements.

Finally, a recent study aimed to identify physiological factors that contribute to the large observed between-subject and between-session variability seen within BOLD-CVR data (Hou et al., 2020); the authors reported a significant negative association between BOLD-CVR and baseline P_ET_CO_2_, across subjects. The relationship between baseline P_ET_CO_2_ and CVR is likely mediated by baseline CBF, yet when performing exploratory correlation analyses, we did not find significant evidence that bCBF or BOLD-CVR was related to baseline P_ET_CO_2_ in our data (Supplementary Table 4). We interpret these results cautiously considering our much smaller sample size compared to the previous work (Hou et al., 2020). Furthermore, our use of a nasal cannula is valid to sample ΔP_ET_CO_2_ (Kim et al., 1997; Nayak et al., 2019; Yosefy et al., 2004), particularly in a healthy sample, but when estimating absolute baseline P_ET_CO_2_ for any individual this is likely to be underestimated due to the contribution of room air during sampling.

### 4.3. Relevance and implications for fMRI research

A key motivation behind understanding the relationship between bCBF and BOLD-CVR is to facilitate appropriate normalization analysis strategies, as has been done for task-based BOLD fMRI. For example, previous work has corrected task-induced BOLD signals by covarying out bCBF (Krishnamurthy et al., 2020) or CVR (Cohen et al., 2004; Liu et al., 2013b) to account for varying neurovascular coupling or vascular physiology. However, based on the conflicting evidence presented in existing literature and the results we report here, we suggest that the coupling between bCBF and BOLD-CVR needs to be understood better before such normalization strategies become common-place, and more consensus needs to exist in the healthy population literature. Though we report a significant positive correlation between bCBF and BOLD-CVR, they only share a proportion of their variance. For example, GM bCBF and lag-optimized GM BOLD-CVR shared 53% and 61-67% of between-subject variance for BH+REST and CDB+REST data, respectively. For the within-subject spatial analysis, the average variance shared between bCBF and BOLD-CVR was 42% and 33% for BH+REST and CDB+REST data, respectively. Therefore, it will be important to consider both these metrics when studying the cerebrovasculature in healthy populations, whilst modelling their effects in ways that account for shared variance. The validity of fMRI normalization strategies or covariate analyses will depend on how physiologically accurate the estimates of bCBF and BOLD-CVR are. Many factors will affect the accuracy; in this paper and our previous work (Stickland et al., 2021) we have shown evidence for two factors affecting CVR estimates (lag-optimization and using breathing task data vs. resting-state data only).

Studying the relationship between bCBF and BOLD/CBF-CVR becomes more interesting, yet more complicated, in situations where they may become uncoupled. In populations where we know or suspect bCBF or CVR to be altered, causing an uncoupling of neuronal activation and blood flow (Para et al., 2017), it is even more important to understand the covariance between bCBF and CVR. However, normalizing fMRI results using these metrics in situations where they are uncoupled or coupled in atypical ways is problematic, particularly if causality is not clear (i.e., whether an impairment of bCBF drives an impairment of CVR, or vice versa). For example, in children with sickle cell disease there are reports of elevated bCBF but lower BOLD and CBF CVR (Kosinski et al., 2017; Václavů et al., 2019); in cognitively normal older adults, greater aortic stiffness has been related to lower regional bCBF and higher CBF-CVR (Jefferson et al., 2018); in healthy adults with a normal fitness range, there are reports of aerobic fitness being associated with lower bCBF but greater CBF-CVR (Foster et al., 2020). There are many more examples in the literature where bCBF and CVR do not show the positive coupling observed in this study. Ultimately, understanding the relationship between bCBF and BOLD/CBF-CVR estimates in healthy populations, and what factors affect this relationship, will help us better assess the extent to which they are uncoupled in pathology, and better interpret the nature of this uncoupling.

### 4.4. Study limitations

This study is limited in its conclusions by its small sample size. We chose to visualize each subject’s data points to be transparent regarding the full data variance, rather than only reporting group summary measures. We only used a single post label delay for our pCASL scan, which did not allow us to estimate and account for arterial arrival times in bCBF quantification. However, the post-labeling delay in our acquisition was chosen to be long enough for labeled spins to reach the imaged tissue in healthy young brains (Alsop et al., 2015b), and therefore perfusion quantification should be accurate. For the vasoactive stimuli, we modulated P_ET_CO_2_ values with a breathing task or characterized resting-state fluctuations in P_ET_CO_2_, as opposed to delivering gas inhalation challenges. Such gas challenges, particularly when administered by feed-forward gas blending systems (Slessarev et al., 2007), would allow more precise and repeatable P_ET_CO_2_ targeting while keeping P_ET_CO_2_ values more stable. This methodology could be implemented in future work to improve mapping of BOLD-CVR and further probe its relationship with bCBF. A limitation already discussed in Section 4.2.2 is that we are using the BOLD signal contrast to reflect CBF changes. Future work should consider the measurement of CBF-CVR as well as BOLD-CVR, when probing the bCBF-CVR relationship across individuals and brain regions. Although it is possible to measure CBF-CVR with breathing task designs (Solis-Barquero et al., 2021), controlled gas inhalation challenges will greatly facilitate accurate CBF-CVR measurements through repeated arterial spin labeling acquisitions that demand sustained stable P_ET_CO_2_ modulations.

### 4.5. Conclusions

In summary, we correlated two separately acquired metrics of cerebrovascular health, baseline CBF and BOLD-CVR (with BOLD-CVR estimates from 3 separate fMRI acquisitions), in 9 healthy individuals. These metrics showed stronger and more consistent positive correlations when CVR was modelled with breathing task data as opposed to resting-state data only, and when CVR timing was accounted for. The stronger relationship between bCBF and BOLD-CVR suggests more physiologically valid CVR estimates when such methodological elements are incorporated. Future research should investigate the basis of the observed positive relationship between bCBF and BOLD-CVR in larger normative samples and across the lifespan. Understanding how baseline vascular physiology relates to dynamic vascular processes is particularly important in clinical cohorts that present with altered vascular and/or metabolic baselines or impaired cerebrovascular reserve.

## ACKNOWLEDGMENTS

This research was supported by the Eunice Kennedy Shriver National Institute of Child Health and Human Development of the National Institutes of Health under award number K12HD073945. K.Z. was supported by an NIH-funded training program (T32EB025766). S.M. was supported by the European Union’s Horizon 2020 research and innovation program (Marie Skłodowska-Curie grant agreement No. 713673), a fellowship from La Caixa Foundation (ID 100010434, fellowship code LCF/BQ/IN17/11620063) and C.C.G was supported by the Spanish Ministry of Economy and Competitiveness (Ramon y Cajal Fellowship, RYC2017-21845), the Basque Government (BERC 2018-2021 and PIBA_2019_104) and the Spanish Ministry of Science, Innovation and Universities (MICINN; PID2019-105520GB-100).

## CrediT statement

Rachael Stickland: Conceptualization, Methodology, Software, Formal Analysis, Investigation, Data Curation, Writing (OD), Writing (RE), Visualization, Project Administration. Kristina Zvolanek: Methodology, Investigation, Writing (RE). Stefano Moia: Methodology, Writing (RE). César Caballero-Gaudes: Methodology, Writing (RE). Molly Bright: Conceptualization, Methodology, Software, Investigation, Resources, Writing (RE), Supervision, Project Administration, Funding Acquisition. The authors would also like to thank the personnel at the Center for Translation Imaging (Northwestern Radiology) for support with study set-up.

## Abbreviations

(ALFF): amplitude of low-frequency fluctuation
(bCBF): baseline cerebral blood flow
(BOLD): blood oxygenation dependent signal
(BH): breath-holding
(CBF): cerebral blood flow
(CVR): cerebrovascular reactivity
(CDB): cued deep breathing
(FDR): false discovery rate
(fALFF): fractional amplitude of low-frequency fluctuation
(FWHM): full width half maximum
(fMRI): functional magnetic resonance imaging
(GM): gray matter
(lag optimization): Lag-Opt
(no lag optimization): No-Op
(P_ET_CO_2_): partial pressure of end tidal carbon dioxide
(pCASL): pseudo-continuous arterial spin labelling
(RSFA): resting-state fluctuation amplitude

## REFERENCES

Alahmadi, A.A.S., 2021. Effects of different smoothing on global and regional resting functional connectivity. Neuroradiology 63, 99–109. https://doi.org/10.1007/S00234-020-02523-8/FIGURES/5

Alsop, D.C., Detre, J.A., Golay, X., Günther, M., Hendrikse, J., Hernandez-Garcia, L., Lu, H., MacIntosh, B.J., Parkes, L.M., Smits, M., van Osch, M.J.P., Wang, D.J.J., Wong, E.C., Zaharchuk, G., 2015a. Recommended implementation of arterial spin-labeled perfusion MRI for clinical applications: A consensus of the ISMRM perfusion study group and the European consortium for ASL in dementia. Magn. Reson. Med. 73, 102–116. https://doi.org/10.1002/mrm.25197

Alsop, D.C., Detre, J.A., Golay, X., Günther, M., Hendrikse, J., Hernandez-Garcia, L., Lu, H., MacIntosh, B.J., Parkes, L.M., Smits, M., van Osch, M.J.P., Wang, D.J.J., Wong, E.C., Zaharchuk, G., 2015b. Recommended implementation of arterial spin-labeled perfusion MRI for clinical applications: A consensus of the ISMRM perfusion study group and the European consortium for ASL in dementia. Magn. Reson. Med. 73, 102–116. https://doi.org/10.1002/mrm.25197

Battisti-Charbonney, A., Fisher, J., Duffin, J., 2011. The cerebrovascular response to carbon dioxide in humans. Physiol. Soc. J Physiol 589, 12. https://doi.org/10.1113/jphysiol.2011.206052

Bright, M.G., Bulte, D.P., Jezzard, P., Duyn, J.H., 2009. Characterization of regional heterogeneity in cerebrovascular reactivity dynamics using novel hypocapnia task and BOLD fMRI. Neuroimage 48, 166–75. https://doi.org/10.1016/j.neuroimage.2009.05.026

Bright, M.G., Donahue, M.J., Duyn, J.H., Jezzard, P., Bulte, D.P., 2011. The effect of basal vasodilation on hypercapnic and hypocapnic reactivity measured using magnetic resonance imaging. J. Cereb. Blood Flow Metab. 31, 426–38. https://doi.org/10.1038/jcbfm.2010.187

Bright, M.G., Murphy, K., 2013. Reliable quantification of BOLD fMRI cerebrovascular reactivity despite poor breath-hold performance. Neuroimage 83, 559–68. https://doi.org/10.1016/j.neuroimage.2013.07.007

Bright, M.G., Whittaker, J.R., Driver, I.D., Murphy, K., 2020. Vascular physiology drives functional brain networks. Neuroimage 217. https://doi.org/10.1016/j.neuroimage.2020.116907

Brown, G.G., Zorrilla, L.T.E., Georgy, B., Kindermann, S.S., Wong, E.C., Buxton, R.B., 2003. BOLD and perfusion response to finger-thumb apposition after acetazolamide administration: Differential relationship to global perfusion. J. Cereb. Blood Flow Metab. 23, 829–837. https://doi.org/10.1097/01.WCB.0000071887.63724.B2

Chappell, M.A., Groves, A.R., Whitcher, B., Woolrich, M.W., 2009. Variational Bayesian Inference for a Nonlinear Forward Model. IEEE Trans. Signal Process. 57, 223–236. https://doi.org/10.1109/TSP.2008.2005752

Chen, C.M., Yang, H.C., Hsieh, H.H., Liao, T.Y., Huang, Y.C., Peng, S.L., 2021. Characterization of regional differences in cerebral vascular response to breath holding using BOLD fMRI. Int. J. Imaging Syst. Technol. 31, 180–188. https://doi.org/10.1002/IMA.22473

Chen, J.J., Gauthier, C.J., 2021. The Role of Cerebrovascular-Reactivity Mapping in Functional MRI: Calibrated fMRI and Resting-State fMRI. Front. Physiol. 12, 361. https://doi.org/10.3389/FPHYS.2021.657362/BIBTEX

Chu, P.P.W., Golestani, A.M., Kwinta, J.B., Khatamian, Y.B., Chen, J.J., 2018. Characterizing the modulation of resting-state fMRI metrics by baseline physiology. Neuroimage 173, 72–87. https://doi.org/10.1016/j.neuroimage.2018.02.004

Cohen, E.R., Rostrup, E., Sidaros, K., Lund, T.E., Paulson, O.B., Ugurbil, K., Kim, S.G., 2004. Hypercapnic normalization of BOLD fMRI: comparison across field strengths and pulse sequences. Neuroimage 23, 613–624. https://doi.org/10.1016/J.NEUROIMAGE.2004.06.021

Cohen, E.R., Ugurbil, K., Kim, S.G., 2002. Effect of basal conditions on the magnitude and dynamics of the blood oxygenation level-dependent fMRI response, Journal of Cerebral Blood Flow and Metabolism. https://doi.org/10.1097/00004647-200209000-00002

Cox, R.W., 1996. AFNI: Software for analysis and visualization of functional magnetic resonance neuroimages. Comput. Biomed. Res. 29, 162–173. https://doi.org/10.1006/cbmr.1996.0014

Cumming, P., Eickhoff, S., Johansen-berg, H., Pike, B., Leahy, R., Liu, T.T., Doherty, J.O., Hickok, G., Beaulieu, C., Beckman, C., Booth, J.R., Brookes, M., Cohen, M.S.M.X., Greenwood, P., He, B., Lerch, J., Lindquist, M., Lowe, M.J., Miller, K.L., Schwartz, S., Bandettini, P.A., Thulborn, K.R., Fox, P.T., Koretsky, A.P., Ogawa, S., Kwong, K.K., Uğurbil, K., Bandettini, P.A., Turner, R., Blamire, A.M., Talavage, T.M., Hall, D.A., Jezzard, P., Cohen, M.S.M.X., Schmitt, F., Wong, E.C., Posse, S., Kim, S.G., van Gelderen, P., Duyn, J.H., Ramsey, N.F., Liu, G., Moonen, C.T.W., Weiskopf, N., Hennig, J., Lin, F.H., Tsai, K.W.K., Chu, Y.H., Witzel, T., Nummenmaa, A., Raij, T., Ahveninen, J., Kuo, W.J., Belliveau, J.W., Glover, G.H., Miller, K.L., Feinberg, D.A., Yacoub, E., Uğurbil, K., Lu, H., van Zijl, P.C.M., Cox, R.W., Goebel, R., Van Essen, David C., Aguirre, G.K., Saad, Z.S., Reynolds, R.C., Fischl, B., Jenkinson, M., Beckmann, C.F., Behrens, T.E.J., Woolrich, M.W., Smith, S.M., Ashburner, J., Woolrich, M.W., Nichols, T.E., Price, C.J., Hyde, J.S., Jesmanowicz, A., Haxby, J. V., Stephan, K.E., Roebroeck, A., Birn, R.M., Poline, J.B., Brett, M., Sporns, O., McIntosh, A.R., Beckmann, C.F., Snyder, A.Z., Raichle, M.E., Evans, A.C., Janke, A.L., Collins, D.L., Baillet, S., Haacke, E.M., Ye, Y., Biswal, B.B., Vul, E., Pashler, H., Song, A.W., Buxton, R.B., Logothetis, N.K., Menon, R.S., Boynton, G.M., Engel, S.A., Heeger, D.J., Hyder, F., Rothman, D.L., Villringer, A., Mandeville, J.B., Silva, A.C., Weisskoff, R.M., Handwerker, D.A., Gonzalez-Castillo, J., D’Esposito, M., Bandettini, P.A., Harel, N., Cheng, K., Gao, J.H., Liu, H.L., Lauritzen, M., Mathiesen, C., Schaefer, K., Thomsen, K.J., Krueger, G., Granziera, C., Laufs, H., Menon, R.S., Jenkins, B.G., Binder, J.R., van Zijl, P.C.M., Hua, J., Lu, H., Hu, X., Yacoub, E., Norris, D.G., McGonigle, D.J., Singh, K.D., Le Bihan, D., Buckner, R.L., Lowe, M.J., Huettel, S.A., Liu, T.T., Malach, R., Maguire, E.A., Petersen, S.E., Dubis, J.W., Courtney, S.M., Clark, V.P., Engel, S.A., Savoy, R.L., Koretsky, A.P., Poldrack, R.A., Wald, L.L., Friston, K., Pike, G.B., Duyn, J.H., Formisano, E., Kriegeskorte, N., Smith, S.M., Bullmore, E., Hasson, U., Honey, C.J., Aguirre, G.K., Detre, J.A., Meyer-lindenberg, A., Johansen-berg, H., Essen, D C Van, Ugurbil, K., Reichenbach, J.R., Rosen, B.R., Savoy, R.L., 2012. Linear systems analysis of the fMRI signal. Neuroimage 62, 852–855. https://doi.org/10.1016/j.neuroimage.2011.08.056

Deckers, P.T., Bhogal, A.A., Dijsselhof, M.B.J., Faraco, C.C., Liu, P., Lu, H., Donahue, M.J., Siero, J.C.W., 2021. Hemodynamic and metabolic changes during hypercapnia with normoxia and hyperoxia using pCASL and TRUST MRI in healthy adults. J. Cereb. Blood Flow Metab. https://doi.org/10.1177/0271678X211064572

Deng, S., Franklin, C.G., O’Boyle, M., Zhang, W., Heyl, B.L., Jerabek, P.A., Lu, H., Fox, P.T., 2022. Hemodynamic and metabolic correspondence of resting-state voxel-based physiological metrics in healthy adults. Neuroimage 250, 118923. https://doi.org/10.1016/J.NEUROIMAGE.2022.118923

Desikan, R.S., Ségonne, F., Fischl, B., Quinn, B.T., Dickerson, B.C., Blacker, D., Buckner, R.L., Dale, A.M., Maguire, R.P., Hyman, B.T., Albert, M.S., Killiany, R.J., 2006. An automated labeling system for subdividing the human cerebral cortex on MRI scans into gyral based regions of interest. Neuroimage 31, 968–980. https://doi.org/10.1016/J.NEUROIMAGE.2006.01.021

Driver, I.D., Wise, R.G., Murphy, K., 2017. Graded hypercapnia-calibrated BOLD: Beyond the Iso-metabolic hypercapnic assumption. Front. Neurosci. 11, 276. https://doi.org/10.3389/fnins.2017.00276

Foster, C., Steventon, J.J., Helme, D., Tomassini, V., Wise, R.G., 2020. Assessment of the Effects of Aerobic Fitness on Cerebrovascular Function in Young Adults Using Multiple Inversion Time Arterial Spin Labeling MRI. Front. Physiol. 11, 360. https://doi.org/10.3389/FPHYS.2020.00360/BIBTEX

Frossard, J., Renaud, O., 2019. permuco: Permutation Tests for Regression, (Repeated Measures) ANOVA/ANCOVA and Comparison of Signals. R package version 1.1.0.

Golestani, A.M., Wei, L.L., Chen, J.J., 2016. Quantitative mapping of cerebrovascular reactivity using resting-state BOLD fMRI: Validation in healthy adults. Neuroimage 138, 147–163. https://doi.org/10.1016/j.neuroimage.2016.05.025

Grabner, G., Janke, A.L., Budge, M.M., Smith, D., Pruessner, J., Collins, D.L., 2006. Symmetric Atlasing and Model Based Segmentation: An Application to the Hippocampus in Older Adults. Lect. Notes Comput. Sci. (including Subser. Lect. Notes Artif. Intell. Lect. Notes Bioinformatics) 4191 LNCS-II, 58–66. https://doi.org/10.1007/11866763_8

Griffeth, V.E.M., Perthen, J.E., Buxton, R.B., 2011. Prospects for quantitative fMRI: Investigating the effects of caffeine on baseline oxygen metabolism and the response to a visual stimulus in humans. Neuroimage 57, 809–816. https://doi.org/10.1016/j.neuroimage.2011.04.064

Halani, S., Kwinta, J.B., Golestani, A.M., Khatamian, Y.B., Chen, J.J., 2015. Comparing cerebrovascular reactivity measured using BOLD and cerebral blood flow MRI: The effect of basal vascular tension on vasodilatory and vasoconstrictive reactivity. Neuroimage 110, 110–123. https://doi.org/10.1016/j.neuroimage.2015.01.050

Hou, X., Liu, P., Li, Y., Jiang, D., De Vis, J.B., Lin, Z., Sur, S., Baker, Z., Mao, D., Ravi, H., Rodrigue, K., Albert, M., Park, D.C., Lu, H., 2020. The association between BOLD-based cerebrovascular reactivity (CVR) and end-tidal CO2 in healthy subjects. Neuroimage 207, 116365. https://doi.org/10.1016/j.neuroimage.2019.116365

Ito, H., Kanno, I., Ibaraki, M., Suhara, T., Miura, S., 2008. Relationship between baseline cerebral blood flow and vascular responses to changes in PaCO2 measured by positron emission tomography in humans: implication of inter-individual variations of cerebral vascular tone. Acta Physiol. 193, 325–330. https://doi.org/10.1111/J.1748-1716.2008.01847.X

Jefferson, A.L., Cambronero, F.E., Liu, D., Moore, E.E., Neal, J.E., Terry, J.G., Nair, S., Pechman, K.R., Rane, S., Davis, L.T., Gifford, K.A., Hohman, T.J., Bell, S.P., Wang, T.J., Beckman, J.A., Carr, J.J., 2018. Higher aortic stiffness is related to lower cerebral blood flow and preserved cerebrovascular reactivity in older adults. Circulation 138, 1951–1962. https://doi.org/10.1161/CIRCULATIONAHA.118.032410

Jenkinson, M., Bannister, P., Brady, M., Smith, S., 2002. Improved Optimization for the Robust and Accurate Linear Registration and Motion Correction of Brain Images. Neuroimage 17, 825–841. https://doi.org/10.1006/nimg.2002.1132

Jenkinson, M., Beckmann, C.F., Behrens, T.E.J., Woolrich, M.W., Smith, S.M., 2012. FSL. Neuroimage 62, 782–790. https://doi.org/10.1016/J.NEUROIMAGE.2011.09.015

Jenkinson, M., Smith, S., 2001. A global optimisation method for robust affine registration of brain images. Med. Image Anal. 5, 143–156. https://doi.org/10.1016/S1361-8415(01)00036-6

Kannurpatti, S.S., Biswal, B.B., 2008. Detection and scaling of task-induced fMRI-BOLD response using resting state fluctuations. Neuroimage 40, 1567–1574. https://doi.org/10.1016/j.neuroimage.2007.09.040

Kannurpatti, S.S., Motes, M.A., Rypma, B., Biswal, B.B., 2010. Neural and vascular variability and the fMRI-BOLD response in normal aging. Magn. Reson. Imaging 28, 466. https://doi.org/10.1016/J.MRI.2009.12.007

Kassambara, A., 2020. ggpubr: “ggplot2” Based Publication Ready Plots. R package version 0.4.0.

Kim, S.G., Ogawa, S., 2012. Biophysical and physiological origins of blood oxygenation level-dependent fMRI signals. J. Cereb. Blood Flow Metab. https://doi.org/10.1038/jcbfm.2012.23

Kim, Y.J., Lee, D.G., Chung, S.H., Choe, Y.K., Park, J.W., Shin, C.M., Park, J.Y., 1997. The Monitoring of PETCO2 via Nasal Cannula in Spontaneously Breathing Patients during Spinal Anesthesia. Korean J. Anesthesiol. 33, 243. https://doi.org/10.4097/KJAE.1997.33.2.243

Kosinski, P.D., Croal, P.L., Leung, J., Williams, S., Odame, I., Hare, G.M.T., Shroff, M., Kassner, A., 2017. The severity of anaemia depletes cerebrovascular dilatory reserve in children with sickle cell disease: a quantitative magnetic resonance imaging study. Br. J. Haematol. 176, 280–287. https://doi.org/10.1111/bjh.14424

Krishnamurthy, V., Krishnamurthy, L.C., Drucker, J.H., Kundu, S., Ji, B., Hortman, K., Roberts, S.R., Mammino, K., Tran, S.M., Gopinath, K., McGregor, K.M., Rodriguez, A.D., Qiu, D., Crosson, B., Nocera, J.R., 2020. Correcting Task fMRI Signals for Variability in Baseline CBF Improves BOLD-Behavior Relationships: A Feasibility Study in an Aging Model. Front. Neurosci. 14, 336. https://doi.org/10.3389/FNINS.2020.00336/BIBTEX

Leoni, R.F., Oliveira, I.A.F., Pontes-Neto, O.M., Santos, A.C., Leite, J.P., 2017. Cerebral blood flow and vasoreactivity in aging: An arterial spin labeling study. Brazilian J. Med. Biol. Res. 50. https://doi.org/10.1590/1414-431X20175670

Leung, J., Kosinski, P.D., Croal, P.L., Kassner, A., 2016. Developmental trajectories of cerebrovascular reactivity in healthy children and young adults assessed with magnetic resonance imaging. J. Physiol. 594, 2681–9. https://doi.org/10.1113/JP271056

Li, X., Morgan, P.S., Ashburner, J., Smith, J., Rorden, C., 2016. The first step for neuroimaging data analysis: DICOM to NIfTI conversion. J. Neurosci. Methods 264, 47–56. https://doi.org/10.1016/j.jneumeth.2016.03.001

Liu, P., Calhoun, V., Chen, Z., 2017. Functional overestimation due to spatial smoothing of fMRI data. J. Neurosci. Methods 291, 1–12. https://doi.org/10.1016/J.JNEUMETH.2017.08.003

Liu, P., Hebrank, A.C., Rodrigue, K.M., Kennedy, K.M., Park, D.C., Lu, H., 2013a. A comparison of physiologic modulators of fMRI signals. Hum. Brain Mapp. 34, 2078–2088. https://doi.org/10.1002/HBM.22053

Liu, P., Hebrank, A.C., Rodrigue, K.M., Kennedy, K.M., Section, J., Park, D.C., Lu, H., 2013b. Age-related differences in memory-encoding fMRI responses after accounting for decline in vascular reactivity. Neuroimage 78, 415. https://doi.org/10.1016/J.NEUROIMAGE.2013.04.053

Liu, T.T., 2013. Neurovascular factors in resting-state functional MRI. Neuroimage 80, 339–348. https://doi.org/10.1016/j.neuroimage.2013.04.071

Lu, H., Xu, F., Rodrigue, K.M., Kennedy, K.M., Cheng, Y., Flicker, B., Hebrank, A.C., Uh, J., Park, D.C., 2011. Alterations in Cerebral Metabolic Rate and Blood Supply across the Adult Lifespan. Cereb. Cortex (New York, NY) 21, 1426. https://doi.org/10.1093/CERCOR/BHQ224

Lu, H., Zhao, C., Ge, Y., Lewis-Amezcua, K., 2008. Baseline blood oxygenation modulates response amplitude: Physiologic basis for intersubject variations in functional MRI signals. Magn. Reson. Med. 60, 364–72. https://doi.org/10.1002/mrm.21686

Marstrand, J.R., Garde, E., Rostrup, E., Ring, P., Rosenbaum, S., Mortensen, E.L., Larsson, H.B.W., 2002. Cerebral perfusion and cerebrovascular reactivity are reduced in white matter hyperintensities. Stroke 33, 972–976. https://doi.org/10.1161/01.STR.0000012808.81667.4B

McSwain, S.D., Hamel, D.S., Smith, P.B., Gentile, M.A., Srinivasan, S., Meliones, J.N., Cheifetz, I.M., 2010. End-tidal and arterial carbon dioxide measurements correlate across all levels of physiologic dead space. Respir. Care 55, 288–93.

Meng, L., Gelb, A.W., 2015. Regulation of cerebral autoregulation by carbon dioxide. Anesthesiology. https://doi.org/10.1097/ALN.0000000000000506

Moia, S., Stickland, R.C., Ayyagari, A., Termenon, M., Caballero-Gaudes, C., Bright, M.G., 2020. Voxelwise optimization of hemodynamic lags to improve regional CVR estimates in breath-hold fMRI, in: Proceedings of the Annual International Conference of the IEEE Engineering in Medicine and Biology Society, EMBS. IEEE, pp. 1489–1492. https://doi.org/10.1109/EMBC44109.2020.9176225

Nayak, V.R., Skariah, A.A., Lewis, T., 2019. Standardization and Validation of Non-invasive Monitoring of End Tidal Carbon Dioxide in Neonates via Nasal Cannula: An Observational Study. Iran. J. Neonatol. 10. https://doi.org/10.22038/ijn.2018.32704.1460

Pajula, J., Tohka, J., 2014. Effects of spatial smoothing on inter-subject correlation based analysis of FMRI. Magn. Reson. Imaging 32, 1114–1124. https://doi.org/10.1016/J.MRI.2014.06.001

Para, A.E., Sam, K., Poublanc, J., Fisher, J.A., Crawley, A.P., Mikulis, D.J., 2017. Invalidation of fMRI experiments secondary to neurovascular uncoupling in patients with cerebrovascular disease. J. Magn. Reson. Imaging 46, 1448–1455. https://doi.org/10.1002/JMRI.25639

Pinto, J., Bright, M.G., Bulte, D.P., Figueiredo, P., 2021. Cerebrovascular Reactivity Mapping Without Gas Challenges: A Methodological Guide. Front. Physiol. 11, 1711. https://doi.org/10.3389/fphys.2020.608475

Schaeffer, S., Iadecola, C., 2021. Revisiting the neurovascular unit. Nat. Neurosci. 24, 1198–1209. https://doi.org/10.1038/s41593-021-00904-7

Slessarev, M., Han, J., Mardimae, A., Prisman, E., Preiss, D., Volgyesi, G., Ansel, C., Duffin, J., Fisher, J.A., 2007. Prospective targeting and control of end-tidal CO2 and O2 concentrations. J. Physiol. 581, 1207–1219. https://doi.org/10.1113/jphysiol.2007.129395

Smith, S.M., 2002. Fast robust automated brain extraction. Hum. Brain Mapp. 17, 143–155. https://doi.org/10.1002/hbm.10062

Smith, S.M., Jenkinson, M., Woolrich, M.W., Beckmann, C.F., Behrens, T.E.J., Johansen-Berg, H., Bannister, P.R., De Luca, M., Drobnjak, I., Flitney, D.E., Niazy, R.K., Saunders, J., Vickers, J., Zhang, Y., De Stefano, N., Brady, J.M., Matthews, P.M., 2004. Advances in functional and structural MR image analysis and implementation as FSL, in: NeuroImage. Academic Press, pp. S208–S219. https://doi.org/10.1016/j.neuroimage.2004.07.051

Sobczyk, O., Battisti-Charbonney, A., Fierstra, J., Mandell, D.M., Poublanc, J., Crawley, A.P., Mikulis, D.J., Duffin, J., Fisher, J.A., 2014. A conceptual model for CO2-induced redistribution of cerebral blood flow with experimental confirmation using BOLD MRI. Neuroimage 92, 56–68. https://doi.org/10.1016/J.NEUROIMAGE.2014.01.051

Solis-Barquero, S.M., Echeverria-Chasco, R., Calvo-Imirizaldu, M., Cacho-Asenjo, E., Martinez-Simon, A., Vidorreta, M., Dominguez, P.D., García de Eulate, R., Fernandez-Martinez, M., Fernández-Seara, M.A., 2021. Breath-Hold Induced Cerebrovascular Reactivity Measurements Using Optimized Pseudocontinuous Arterial Spin Labeling. Front. Physiol. 12, 138. https://doi.org/10.3389/FPHYS.2021.621720/BIBTEX

Sousa, I., Vilela, P., Figueiredo, P., 2014. Reproducibility of hypocapnic cerebrovascular reactivity measurements using BOLD fMRI in combination with a paced deep breathing task. Neuroimage 98, 31–41. https://doi.org/10.1016/j.neuroimage.2014.04.049

Stefanovic, B., Warnking, J.M., Rylander, K.M., Pike, G.B., 2006. The effect of global cerebral vasodilation on focal activation hemodynamics. Neuroimage 30, 726–734. https://doi.org/10.1016/J.NEUROIMAGE.2005.10.038

Stickland, R.C., Zvolanek, K.M., Moia, S., Ayyagari, A., Caballero-Gaudes, C., Bright, M.G., 2021. A practical modification to a resting state fMRI protocol for improved characterization of cerebrovascular function. Neuroimage 239, 118306. https://doi.org/10.1016/J.NEUROIMAGE.2021.118306

Suri, S., Mackay, C.E., Kelly, M.E., Germuska, M., Tunbridge, E.M., Frisoni, G.B., Matthews, P.M., Ebmeier, K.P., Bulte, D.P., Filippini, N., 2015. Reduced cerebrovascular reactivity in young adults carrying the APOE ε4 allele. Alzheimer’s Dement. 11, 648–657.e1. https://doi.org/10.1016/J.JALZ.2014.05.1755

Tak, S., Wang, D.J.J., Polimeni, J.R., Yan, L., Chen, J.J., 2014. Dynamic and static contributions of the cerebrovasculature to the resting-state BOLD signal. Neuroimage 84, 672–680. https://doi.org/10.1016/j.neuroimage.2013.09.057

Takano, Y., Sakamoto, O., Kiyofuji, C., Ito, K., 2003. A comparison of the end-tidal CO2 measured by portable capnometer and the arterial PCO2 in spontaneously breathing patients, Respiratory Medicine. https://doi.org/10.1053/rmed.2002.1468

Tancredi, F.B., Hoge, R.D., 2013. Comparison of cerebral vascular reactivity measures obtained using breath-holding and CO2 inhalation. J. Cereb. Blood Flow Metab. 33, 1066–74. https://doi.org/10.1038/jcbfm.2013.48

Tisdall, M.D., Reuter, M., Qureshi, A., Buckner, R.L., Fischl, B., van der Kouwe, A.J.W., 2016. Prospective motion correction with volumetric navigators (vNavs) reduces the bias and variance in brain morphometry induced by subject motion. Neuroimage 127, 11–22. https://doi.org/10.1016/j.neuroimage.2015.11.054

Tsvetanov, K.A., Henson, R.N.A., Rowe, J.B., 2021. Separating vascular and neuronal effects of age on fMRI BOLD signals. Philos. Trans. R. Soc. B 376, 20190631. https://doi.org/10.1098/RSTB.2019.0631

Urback, A.L., MacIntosh, B.J., Goldstein, B.I., 2017. Cerebrovascular reactivity measured by functional magnetic resonance imaging during breath-hold challenge: A systematic review. Neurosci. Biobehav. Rev. https://doi.org/10.1016/j.neubiorev.2017.05.003

Václavů, L., Meynart, B.N., Mutsaerts, H.J.M.M., Petersen, E.T., Majoie, C.B.L.M., Vanbavel, E.T., Wood, J.C., Nederveen, A.J., Biemond, B.J., 2019. Hemodynamic provocation with acetazolamide shows impaired cerebrovascular reserve in adults with sickle cell disease. Haematologica 104, 690. https://doi.org/10.3324/HAEMATOL.2018.206094

Vazquez, A.L., Cohen, E.R., Gulani, V., Hernandez-Garcia, L., Zheng, Y., Lee, G.R., Kim, S.G., Grotberg, J.B., Noll, D.C., 2006. Vascular dynamics and BOLD fMRI: CBF level effects and analysis considerations. Neuroimage 32, 1642–1655. https://doi.org/10.1016/J.NEUROIMAGE.2006.04.195

Vigneau-Roy, N., Bernier, M., Descoteaux, M., Whittingstall, K., 2014. Regional variations in vascular density correlate with resting-state and task-evoked blood oxygen level-dependent signal amplitude. Hum. Brain Mapp. 35, 1906. https://doi.org/10.1002/HBM.22301

Wickham, H., 2016. ggplot2: Elegant Graphics for Data Analysis.

Woolrich, M.W., Jbabdi, S., Patenaude, B., Chappell, M., Makni, S., Behrens, T., Beckmann, C., Jenkinson, M., Smith, S.M., 2009. Bayesian analysis of neuroimaging data in FSL. Neuroimage 45, S173–S186. https://doi.org/10.1016/j.neuroimage.2008.10.055

Xu, F., Uh, J., Brier, M.R., Hart, J., Yezhuvath, U.S., Gu, H., Yang, Y., Lu, H., 2011. The influence of carbon dioxide on brain activity and metabolism in conscious humans. J. Cereb. Blood Flow Metab. 31, 58–67. https://doi.org/10.1038/jcbfm.2010.153

Yablonskiy, D.A., 2011. Cerebral metabolic rate in hypercapnia: Controversy continues. J. Cereb. Blood Flow Metab. 31, 1502–1503. https://doi.org/10.1038/jcbfm.2011.32

Yosefy, C., Hay, E., Nasri, Y., Magen, E., Reisin, L., 2004. End tidal carbon dioxide as a predictor of the arterial PCO 2 in the emergency department setting. Emerg Med J 21, 557–559. https://doi.org/10.1136/emj.2003.005819

Zang, Y.F., Yong, H., Chao-Zhe, Z., Qing-Jiu, C., Man-Qiu, S., Meng, L., Li-Xia, T., Tian-Zi, J., Yu-Feng, W., 2007. Altered baseline brain activity in children with ADHD revealed by resting-state functional MRI. Brain Dev. 29, 83–91. https://doi.org/10.1016/J.BRAINDEV.2006.07.002

Zhang, Y., Brady, M., Smith, S., 2001. Segmentation of brain MR images through a hidden Markov random field model and the expectation-maximization algorithm. IEEE Trans. Med. Imaging 20, 45–57. https://doi.org/10.1109/42.906424

Zhou, Y., Rodgers, Z.B., Kuo, A.H., 2015. Cerebrovascular reactivity measured with arterial spin labeling and blood oxygen level dependent techniques. Magn. Reson. Imaging 33, 566–576. https://doi.org/10.1016/j.mri.2015.02.018

Zou, Q.H., Zhu, C.Z., Yang, Y., Zuo, X.N., Long, X.Y., Cao, Q.J., Wang, Y.F., Zang, Y.F., 2008. An improved approach to detection of amplitude of low-frequency fluctuation (ALFF) for resting-state fMRI: fractional ALFF. J. Neurosci. Methods 172, 137–141. https://doi.org/10.1016/J.JNEUMETH.2008.04.012

