## Supplementary Data for "Lag-optimized BOLD cerebrovascular reactivity derived from breathing task data has a stronger relationship with baseline cerebral blood flow"

### SUPPLEMENTARY MATERIAL

| **GM Mask** and justification (pv= partial volume) | **Data Segment (Opt/NoOpt)** | **r** | **t** | **p-val** | **ID of outlier subject(s)** | **r** | **t** | **p-val** |
|---|---|---|---|---|---|---|---|---|
| **Pv threshold = 0.5.** Equal voxel numbers for all data segments. Largest representation of GM voxels, compared to other masks. | BH+R (Opt) | 0.73 | 2.8 | ***0.0264*** | | | | |
| | BH+R (NoOpt) | 0.33 | 0.95 | 0.3782 | | | | |
| | CDB+R(Opt) | 0.78 | 3.35 | ***0.0123*** | 6 | 0.82 | 3.53 | ***0.0123*** |
| | CDB+R (NoOpt) | 0.6 | 1.99 | 0.0864 | 6 | 0.61 | 1.89 | 0.1082 |
| | REST (Opt) | 0.39 | 1.12 | 0.2992 | | | | |
| | REST (NoOpt) | -0.23 | -0.62 | 0.5579 | 6 | -0.52 | -1.51 | 0.1819 |
| | $REST_{BH}$ (Opt) | 0.52 | 1.62 | 0.149 | | | | |
| | $REST_{BH}$ (NoOpt) | 0.38 | 1.1 | 0.3089 | | | | |
| | $REST_{CDB}$ (Opt) | -0.03 | -0.08 | 0.9372 | 6 | 0.45 | 1.24 | 0.2627 |
| | $REST_{CDB}$ (NoOpt) | 0.24 | 0.66 | 0.5302 | 3 and 9 | 0.37 | 0.9 | 0.4092 |
| **Pv threshold = 0.5, no voxels at lag boundaries.** Lag-Opt CVR values at boundary are less accurate. However, different number of voxels included across segments. | BH+R (Opt) | 0.68 | 2.45 | ***0.0438*** | | | | |
| | BH+R (NoOpt) | 0.31 | 0.85 | 0.423 | | | | |
| | CDB+R(Opt) | 0.7 | 2.6 | ***0.0353*** | 6 | 0.76 | 2.87 | ***0.0283*** |
| | CDB+R (NoOpt) | 0.55 | 1.75 | 0.1231 | 6 | 0.53 | 1.54 | 0.1756 |
| | REST (Opt) | 0.37 | 1.04 | 0.333 | 9 | 0.45 | 1.24 | 0.2604 |
| | REST (NoOpt) | -0.34 | -0.96 | 0.3677 | 6 | -0.62 | -1.96 | 0.0983 |
| | $REST_{BH}$ (Opt) | 0.5 | 1.51 | 0.1749 | | | | |
| | $REST_{BH}$ (NoOpt) | 0.29 | 0.8 | 0.4482 | | | | |
| | $REST_{CDB}$ (Opt) | -0.16 | -0.42 | 0.6872 | 6 | 0.24 | 0.61 | 0.5624 |
| | $REST_{CDB}$ (NoOpt) | 0.06 | 0.15 | 0.8848 | 3 and 9 | 0.07 | 0.15 | 0.8875 |
| **Pv threshold = 0.75.** Higher threshold reduces pv effects, whilst maintaining the equal voxel numbers across data segments. | BH+R (Opt) | 0.72 | 2.78 | ***0.0273*** | | | | |
| | BH+R (NoOpt) | 0.34 | 0.97 | 0.3662 | | | | |
| | CDB+R(Opt) | 0.78 | 3.3 | ***0.0131*** | 6 | 0.81 | 3.43 | ***0.014*** |
| | CDB+R (NoOpt) | 0.61 | 2.04 | 0.0802 | 6 | 0.63 | 1.97 | 0.0966 |
| | REST (Opt) | 0.18 | 0.49 | 0.6418 | 7 | 0.56 | 1.66 | 0.1487 |
| | REST (NoOpt) | -0.19 | -0.51 | 0.6254 | 6 | -0.5 | -1.41 | 0.209 |
| | $REST_{BH}$ (Opt) | 0.53 | 1.65 | 0.1439 | | | | |
| | $REST_{BH}$ (NoOpt) | 0.4 | 1.14 | 0.2904 | | | | |
| | $REST_{CDB}$ (Opt) | -0.02 | -0.06 | 0.9528 | 6 | 0.47 | 1.29 | 0.2454 |
| | $REST_{CDB}$ (NoOpt) | 0.25 | 0.69 | 0.5141 | 3 and 9 | 0.43 | 1.07 | 0.3328 |

**Supplementary Table 1.** Correlation results between median GM CBF values and median GM BOLD-CVR values, shown for each analysis and masking option. CVR values across subjects were normally distributed for each test. P-values are not corrected for multiple comparisons.

| Contrast | Mean Diff | t | q |
|---|---|---|---|
| BH+REST - CDB+REST | 0.1219 | 1.362 | 0.3045 |
| BH+REST - REST | 0.1115 | 1.246 | 0.3169 |
| BH+REST - $REST_{BH}$ | 0.1868 | 2.086 | 0.1125 |
| BH+REST - $REST_{CDB}$ | 0.3381 | 3.777 | 0.0065 |
| CDB+REST - REST | -0.0104 | -0.116 | 0.9083 |
| CDB+REST - $REST_{BH}$ | 0.0648 | 0.724 | 0.5267 |
| CDB+REST - $REST_{CDB}$ | 0.2161 | 2.415 | 0.0722 |
| REST - $REST_{BH}$ | 0.0752 | 0.841 | 0.5086 |
| REST - $REST_{CDB}$ | 0.2265 | 2.531 | 0.0722 |
| $REST_{BH}$ - $REST_{CDB}$ | 0.1513 | 1.69 | 0.2014 |

**Supplementary Table 2.** Pairwise comparisons between the five data segments, explored after a significant main effect of data segment was found. Differences are of Fisher's Z values, representing the spatial relationship between bCBF and BOLD-CVR values, presented in Figure 4 of the results. As there was no interaction effect, these mean differences are averaged over each level of optimization (no optimization, lag optimization). q = FDR corrected p-values.

| Sub-ID | Opt/NoOpt | BH+REST | CDB+REST | REST | $REST_{BH}$ | $REST_{CDB}$ |
|---|---|---|---|---|---|---|
| **1** | NoOpt | 8 | 7 | 8 | 5 | 3 |
| **2** | NoOpt | 4 | 3 | 7 | 7 | 4 |
| **3** | NoOpt | 3 | 6 | 3 | 5 | 4 |
| **4** | NoOpt | 3 | 4 | 5 | 3 | 5 |
| **5** | NoOpt | 7 | 5 | 8 | 6 | 5 |
| **6** | NoOpt | 5 | 7 | 3 | 4 | 6 |
| **7** | NoOpt | 6 | 6 | 5 | 3 | 5 |
| **8** | NoOpt | 6 | 4 | 9 | 3 | 5 |
| **9** | NoOpt | 5 | 6 | 6 | 4 | 3 |
| **1** | Opt | 6 | 7 | 12 | 5 | 8 |
| **2** | Opt | 7 | 4 | 3 | 8 | 7 |
| **3** | Opt | 3 | 7 | 1 | 7 | 5 |
| **4** | Opt | 4 | 4 | 5 | 1 | 4 |
| **5** | Opt | 8 | 8 | 8 | 9 | 4 |
| **6** | Opt | 6 | 7 | 8 | 4 | 7 |
| **7** | Opt | 7 | 4 | 6 | 3 | 3 |
| **8** | Opt | 7 | 3 | 5 | 8 | 6 |
| **9** | Opt | 3 | 6 | 3 | 4 | 3 |
| | ***NoOpt Mean*** | *5.22* | *5.33* | *6.00* | *4.44* | *4.44* |
| | ***Opt Mean*** | *5.67* | *5.56* | *5.67* | *5.44* | *5.22* |

**Supplementary Table 3.** The number of parcels removed (out of a total of 96) from the spatial correlation analyses presented in Figure 4, for each subject and analysis type. Parcels were removed when classed as influential points based on Cook's distance.

| **A.** | *Section 1* | | |
|---|---|---|---|
| *Section 2* | $r_s(7)$=0.88,*p=0.0031* | *Section 2* | |
| *Section 3* | $r_s(7)$=0.88,*p=0.0031* | $r_s(7)$=0.93, *p=0.0007* | *Section 3* |
| *Section 4* | $r_s(7)$=1.0,*p<0.00001* | $r_s(7)$=0.88, *p=0.0031* | $r_s(7)$=0.88,*p=0.0031* |

| **B.** | 34.79, 36.23, 36.40, 36.61, 38.65, 39.23, 41.51, 42.32, 42.42 |
|---|---|

| **C.** | *No-Opt* | *Opt* |
|---|---|---|
| *BOLD-CVR, BH+REST* | $r_s(7)$=0.30,*p*=0.4366 | $r_s(7)$=0.33, *p*=0.3853 |
| *BOLD-CVR, CDB+REST* | $r_s(7)$=0.2, *p*=0.6134 | $r_s(7)$=0.25, *p*=0.5206 |
| *BOLD-CVR, REST* | $r_s(7)$=0.15,*p*=0.7000 | $r_s(7)$=-0.07,*p*=0.8801 |
| *BOLD-CVR,* $REST_{BH}$ | $r_s(7)$=0.17,*p*=0.6777 | $r_s(7)$=0.32, *p*=0.4101 |
| *BOLD-CVR,* $REST_{CDB}$ | $r_s(7)$=0.63,*p*=0.0760 | $r_s(7)$=0.83, p=0.0083 |
| *bCBF* | rs(7)=0.17, *p*=0.6777 | |

**Supplementary Table 4. (A)** The relationship between four estimates of baseline $P_{ET}CO_2$, across subjects, averaged over non-overlapping time segments. Section 1 is during the resting pCASL scan (excluding the M0 volume and first 2 data volumes); Section 2 is during the BH+REST fMRI scan (excluding the BH task and the 2 minutes following); Section 3 is during the CDB+REST scan (excluding the CDB task and the 2 minutes following); Section 4 is during the REST scan (excluding the first 2 minutes). **(B)** Average baseline $P_{ET}CO_2$ value, across the 4 sections described in A, for each subject. Due to all four baseline $P_{ET}CO_2$ averages strongly positively correlating (if a subject had a higher baseline $P_{ET}CO_2$ in one scan they tended to have a higher in another), we used one baseline $P_{ET}CO_2$ average to correlate with BOLD-CVR and bCBF values. **(C)** The relationship between the average baseline $P_{ET}CO_2$ value and a BOLD-CVR or bCBF GM average value, across subjects. Average baseline $P_{ET}CO_2$ did not significantly correlate with GM bCBF. Baseline $P_{ET}CO_2$ did not significantly correlate with any GM BOLD-CVR values, except for $REST_{CDB}$ data segment, however this is no longer significant after FDR correction for multiple comparisons. $r_s$ *= spearman's rank correlation coefficient.*

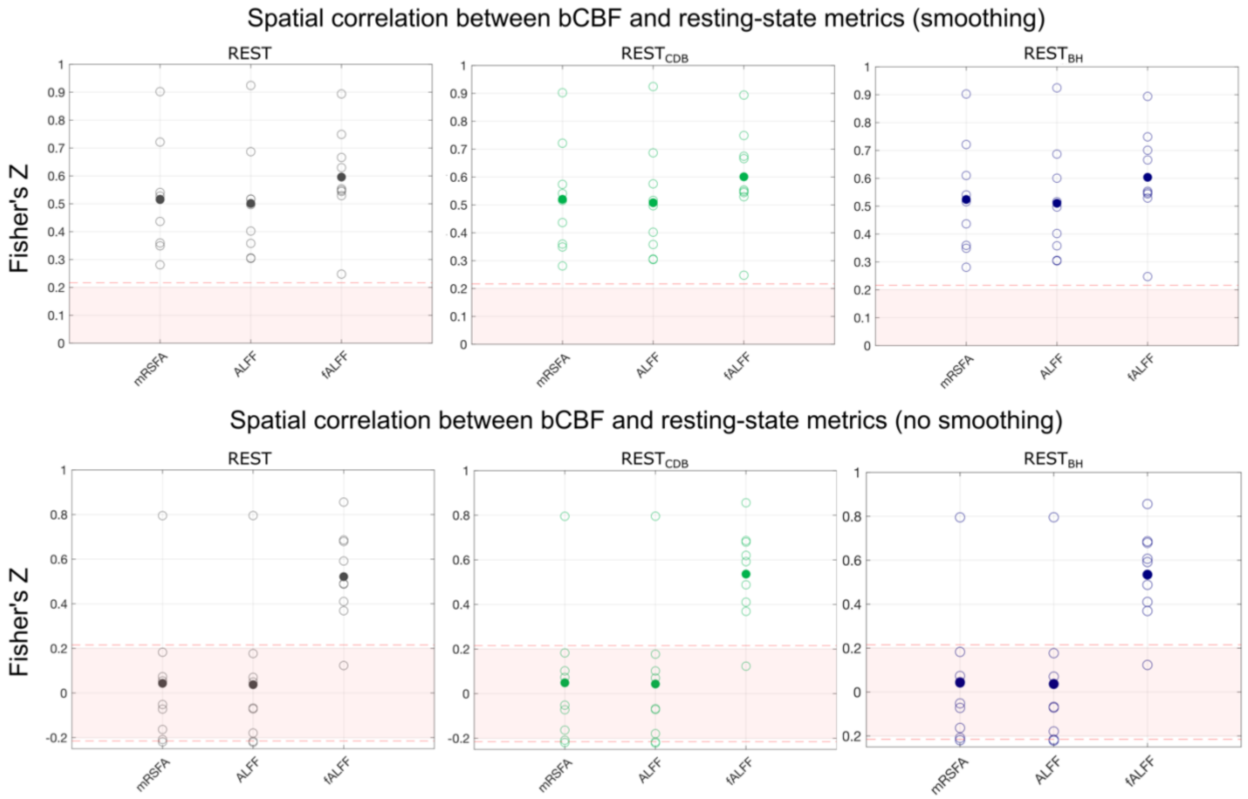


**Supplementary Figure 1.** The association between bCBF and each resting-state metric, in 96 cortical parcels, was computed for each subject. This is shown for the REST segment (column 1), the $REST_{CDB}$ segment (column 2) and the $REST_{BH}$ segment (column 3). Results are shown with (top row) and without (bottom row) spatial smoothing. A Fisher's Z transformation is applied to the correlation values before averaging across subjects. The shaded pink box represents correlation values that would **not be** significant at the single-subject level, based on 96 datapoints and a critical value of 1.96 (p<0.05, two-tailed). When generating these correlation values for each subject, outlier parcels (based on Cook's distance) were not included. As removing parcels reduces the degrees of freedom, the dotted pink line shows the adjusted critical value (maximum across participants and resting-state metric shown for reference).

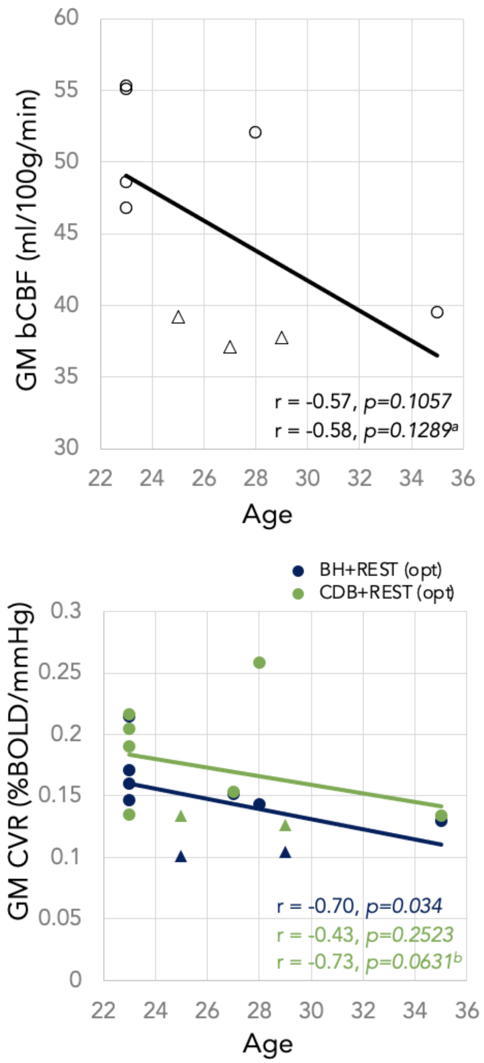


**Supplementary Figure 2.** Top: relationship between Age and GM baseline CBF (bCBF). Bottom: Relationship between Age and GM-CVR for BH+REST (blue) and CDB+REST (green), both lag-optimized ("opt"). Spearman correlation analysis results are shown. [a] 35-year-old participant removed and [b]35 and 28-year-old participant removed as influential points. In both plots, the triangles indicate male participants, and the circles indicate female participants.
